# The genomic basis of copper tolerance in *Drosophila* is shaped by a complex interplay of regulatory and environmental factors

**DOI:** 10.1101/2021.07.12.452058

**Authors:** Llewellyn Green, Marta Coronado-Zamora, Santiago Radio, Gabriel E. Rech, Judit Salces-Ortiz, Josefa González

## Abstract

Increases in industrialization and anthropogenic activity have resulted in an increase of pollutants released into the environment. Of these pollutants, heavy metals such as copper are particularly concerning due to their bio-accumulative nature. Due to its highly heterogeneous distribution and its dual nature as an essential micronutrient and toxic element, the genetic basis of copper tolerance is likely shaped by a complex interplay of genetic and environmental factors.

In this study, we utilized the natural variation present in multiple populations of *Drosophila melanogaster* collected across Europe to screen for variation in copper tolerance. We found that latitude and the degree of urbanization at the collection sites, rather than any other combination of environmental factors, were linked to copper tolerance. While previously identified copper-related genes were not differentially expressed in tolerant *vs*. sensitive strains, genes involved in metabolism, reproduction, and protease induction contributed to the differential stress response. Additionally, the greatest transcriptomic and physiological responses to copper toxicity were seen in the midgut; where we found that preservation of gut acidity is strongly linked to greater tolerance. Finally, we identified transposable element insertions likely to play a role in copper stress response.

Overall, by combining genome-wide approaches with environmental association analysis, and functional analysis of candidate genes, our study provides a unique perspective on the genetic and environmental factors that shape copper tolerance in natural *D. melanogaster* populations, and identifies new genes, transposable elements and physiological traits involved in this complex phenotype.

## BACKGROUND

Rapid industrialization and urbanization have had adverse impacts on biodiversity across ecosystems. Of the contaminants released into the environment due to an increase in human activity, heavy metals are particularly concerning due to their ability to bio-accumulate in soils. Specifically with regard to copper, anthropogenic sources are thought to have a greater influence on topsoil concentrations than either lithological or geographic factors [1]. Human sources of copper are characterized by many point sources of contamination, which has resulted in a highly heterogeneous environmental distribution [2], even across relatively short geographic distances [3]. Due to its highly heterogeneous distribution, and its dual nature as both an essential micronutrient and toxic element, the genetic basis of copper tolerance has the potential to be shaped by a complex interplay of environmental and regulatory factors.

As a commensal species, *Drosophila melanogaster* has a well-documented history as a sentinel of environmental toxins and can be readily sampled from a wide range of geographic locations, making it a prime choice species for the study of copper stress response [4]. *D. melanogaster* has also served as an important tool in the characterization of copper homeostasis and copper-related diseases [5, 6]. As copper acts as an essential micronutrient at low doses but can produce free radicals and damage DNA in excess, the mediation of copper often involves a complex system of regulators, chaperones, and transporters that are commonly found conserved across a wide range of species. Genetic manipulation of *D. melanogaster* has been used to successfully characterize the roles of the common metal-responsive transcription factor-1 (MTF-1) [7], *marvolio* and the *Ctr1* family of transporters which mediate copper uptake [8, 9], the *ATP7* transporter, which regulates copper efflux [10], and the cysteine-rich metallothioneins, which serve to sequester metal ions [11, 12]. Excess copper accumulates in the mid-gut as the fly ages, which is thought to alter gut physiology [13]. Once copper crosses the gut endothelium, it is sequestered by the metallothioneins in the morphologically distinct copper cells and deposited in insoluble granules in the lysozymes [14]. Despite the name, copper cells are considered ‘cuprophobic’ and are inhibited by excess copper [15]. They are also responsible for stomach acid secretion, a function that is lost with age or gut damage, leading to an increase in pH [13].

While many of the aforementioned genes have had their roles in copper homeostasis validated in laboratory conditions, it is not known whether these same genes have an effect on the phenotype in natural populations. To date, there have been several studies exploring the nature of copper tolerance in natural strains of *D. melanogaster*, both with regard to individual genes [16, 17] and to broader developmental and learning and memory processes [18, 19]. Recently, Everman *et al*. (2021) [20] took benefit of a combination of high throughput genomic and transcriptomic approaches to uncover several new copper gene candidates, using recombinant inbred lines. They found that copper resistance is genetically complex and impacted by variation in copper avoidance behavior. In addition to identifying natural variants involved in response to copper, their pairing of genomic data with transcriptomic data also provided a greater opportunity to identify factors that regulate copper induced changes in expression, beyond the well-known MTF-1 factor [20]. Prior expression analyses on metal exposure also suggest that there are a number of co-regulated gene clusters linked to broader stress and metabolism related pathways in response to heavy metal exposure, independent of MTF-1 [20–22]. However, the factors responsible for these coordinated changes in expression have not yet been identified.

To date, genome-wide studies investigating the genetic basis of tolerance to copper and other heavy metals in *D. melanogaster* have focused on SNP variants or were naïve to the nature of the causal variant [20, 23]. The recent availability of new whole-genome assemblies based on long-read sequencing gives us the unprecedented opportunity to characterize complex forms of sequence variation that may have previously been overlooked [24, 25]. This is of particular importance with regard to transposable element insertions, which are often associated with changes in gene expression under stressful conditions (*e*.*g*. [26–31]). Indeed, a natural transposable element insertion in the MTF-1 targeted gene *kuzbanian* has been associated with increased tolerance to zinc in adult flies, although the effect of the insertion was background dependent [32].

In this study, we set out to assess variation in copper tolerance between natural populations of European *D. melanogaster* and investigate whether the phenotype is influenced by either geographic factors, the concentration of copper in soils, atmospheric pollution, or degree of urbanization. To better elucidate the genetic basis of copper tolerance in natural populations, we compared the transcriptomes of three copper tolerant and three copper sensitive strains from before and after copper treatment, using a combination of tissue enrichment analysis, gene ontology, and modular clustering, to examine patterns of gene co-regulation. Finally, we also investigated the physiological traits relevant for copper tolerance. We found that while copper tolerance is highly variable across much of Western Europe, the external factors involved in shaping these phenotypes are complex, likely controlled by multiple regulatory factors, and that tolerance is linked to gut physiology.

## RESULTS

### Copper Tolerance is a Variable Trait Across European *D. melanogaster* Associated with Latitude and Degree of Urbanization

To assess the degree of copper tolerance in natural populations of *D. melanogaster* in Europe, we scored a total of 71 inbred strains, collected from nine locations by the DrosEU consortium (www.droseu.net), for copper mortality on a single dose until full mortality was achieved (Fig. 1A, Table S1A). LT_50_ values ranged from 26.4 to 81.2 hours, with a median value of 49.8 hours (Fig. S1A, Table S2A). We observed very little zero-dose control mortality over the course of the assay (Table S2B). Although we observed a high degree of within-population variance in copper tolerance (Fig. 1B), a linear regression between fly collection locations and LT_50_ values was not significant (p-value = 0.0744) (Table S2D).

**Figure 1.**
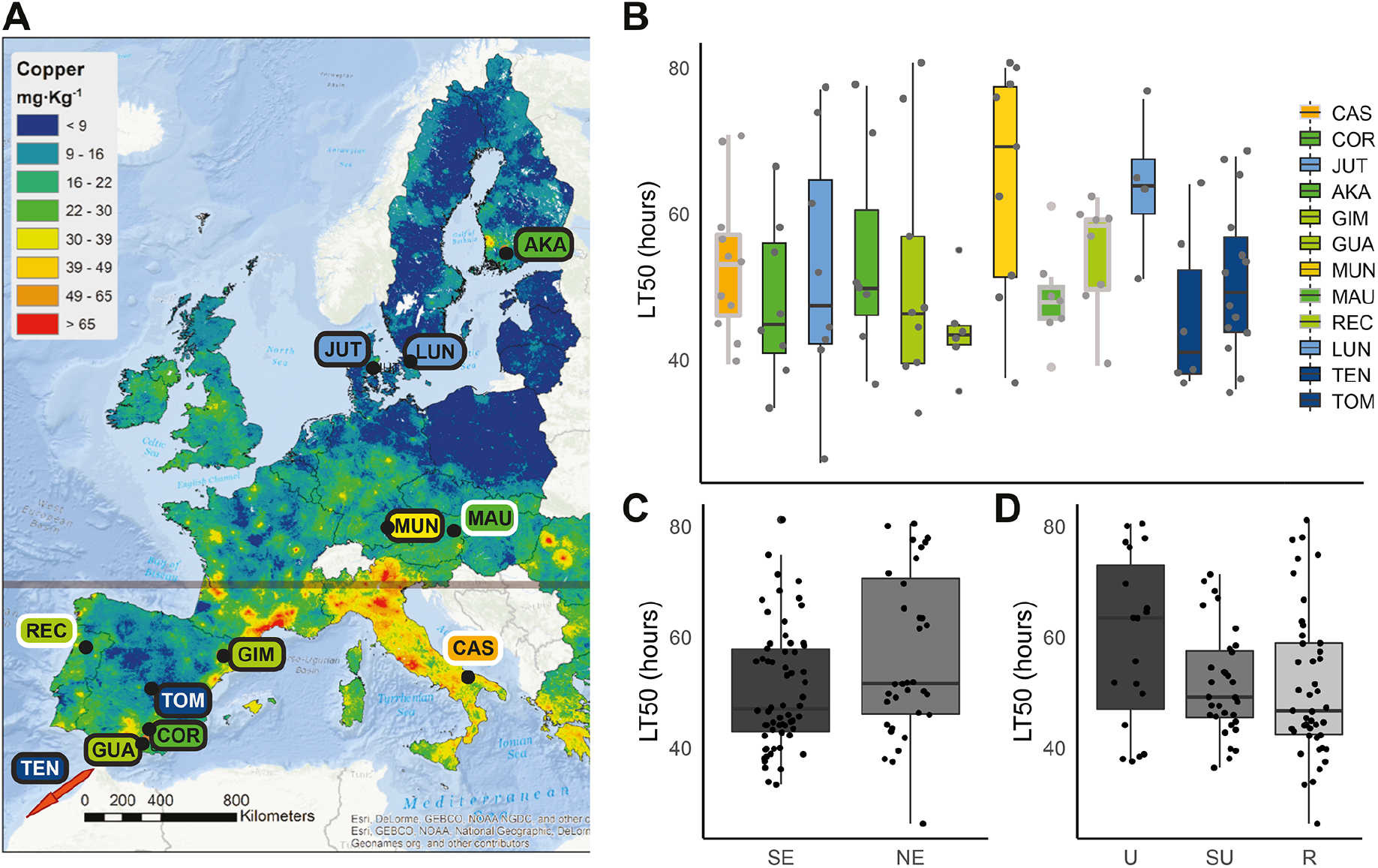
Sampling locations and variation of copper tolerance across Europe. **A)** Distribution of the nine European locations from the 2015 DrosEU collection (black border), and three additional collection locations (white border). Each population is labeled with the first three letters of the collection location (Table S1). Label fill corresponds with the copper concentration legend. The map shows the spatial variation in copper topsoil concentrations, as obtained from the Land Use/Land Cover Area Frame Survey (LUCAS) topsoil database, whose samples were taken from 2009 onwards. The grey line marks the 45th parallel. **B)** Boxplots of LT_50_ values by location. Locations from the 2015 DrosEU collection are outlined in black, while the three additional locations are outlined in grey. The full list of strains used is provided in Table S1A-B. **C)** Boxplots of LT_50_ values of strains split into northern (NE) and southern (SE) European locations by the 45th parallel. This point was chosen because the nine original collection sites could be clearly divided by this line (Fig1A). **D)** Boxplots of LT_50_ values of strains classified by degree of urbanization, U: urban; SU: semi-urban; and R: rural.

As stress tolerance is frequently clinal in *Drosophila* [33–35], we compared the differences in tolerance between northern and southern populations divided by the 45^th^ parallel, as our nine collections sites could be clearly divided by this feature (Fig. 1A). Although the differences in copper tolerance were significant (Wilcoxon’s rank-sum test, p-value = 0.0106), because all southern populations were collected in Spain, we broadened the analysis by phenotyping an additional 19 strains from Portugal and Italy in the south of Europe, along with another 7 strains from Austria (Fig. S1B). As the Portuguese and Austrian strains were caught in 2018 and the Italian strains in 2011, these strains have experienced different degrees of prior laboratory adaptation compared with the previously analyzed nine locations that were all collected in 2015. We found that the association between tolerance and geography was still significant after the inclusion of these new data (p-value of = 0.0378, Fig. 1C).

We further examined the relationship between copper tolerance and geography, copper soil concentrations, atmospheric pollution, and degree of urbanization by fitting a generalized linear model between these potential explanatory variables and the LT50 values across all twelve locations. Longitude was considered in addition to latitude, because European *D. melanogaster* have been reported to exhibit population structure along this axis [36]. As our initial interest in copper was spurred in part by its role as an environmental contaminant, we also tested the relationship between copper tolerance and metal pollution by considering copper concentration in topsoils (mgkg^-1^) and atmospheric pollution (PM10 and PM2.5; general and metal specific), obtained from publicly available data (see Material and Methods; Table S3A). Besides, and as copper contamination is often the result of a complex group of contamination sources, especially around urban areas [2], we also considered a more indirect measure of pollution: degree of urbanization. We classified each of the fly collection locations into urban, semi-urban and rural classes, based on distance from high-, semi- and low-population density areas, ([37]; Table S3A). The final model after performing a backward stepwise regression to eliminate the least significant variables only kept latitude and degree of urbanization (R^2^ = 12%, p-value= 0.0079): we found a positive correlation between latitude and LT_50_ (p-value = 0.015), and we found that urban populations have a higher LT_50_ compared with rural populations (p-value = 0.0086; Table S3B).

### Tolerant and Sensitive Strains Demonstrate Differential Expression Profiles After Copper Exposure Mostly Concentrated in the Midgut

To examine the gene regulatory changes that occur in *D. melanogaster* in response to copper exposure, we compared mated female whole-body transcriptomic profiles of three tolerant (GIM-012, MUN-020, MUN-008) and three sensitive strains (AKA-018, JUT-008 and COR-018), chosen primarily on the basis of their position at the tails of the phenotypic distribution (Fig S1A; see Material and Methods). Carrying out this analysis with strains from the ends of the distribution should be informative about the genes with the greater effects on the phenotypic response. Because we are interested in defining the genes that differentiate the copper tolerant from the copper sensitive strains, we performed DGE analyses for tolerant and sensitive strains separately. Across the three tolerant strains, 239 genes were significantly differentially expressed (> 1.5 fold change and adjusted p-value < 0.05) between copper treatment and control conditions, while 984 genes were differentially expressed across the three sensitive strains, with an overlap of 152 genes (Fig. 2A and Table S4A). Of these 152 genes, the direction of the change was discordant in six genes, being all up-regulated in tolerant strains and down-regulated in sensitive (Table 1). The proportion of down-regulated genes was higher in the sensitive strains, with most of these down-regulated genes unique to sensitive strains (Fig. 2A and Table S4A). These differences in DEGs between tolerant and sensitive strains are also reflected in the Principal component (PC) analysis, where treated and control samples from the JUT-008 and COR-018 showed a much greater separation on PC2 projection (Fig. S2A).

**Table 1.** Differentially Expressed Genes up-regulated in the tolerant and down-regulated in the sensitive strains. ND is no data.

| Gene name | Biological process | Molecular function | Prior copper response association |
| --- | --- | --- | --- |
| *CG13078* | ND | heme binding, oxidoreductase activity, transmembrane ascorbate ferrioreductase activity | Within QTL (Everman et al. 2021) |
| *CG31091* | lipid metabolic process | sterol esterase activity, triglyceride lipase activity | ND |
| *Hml* | hemolymph coagulation, hemostasis, wound healing | protein homodimerization activity, chitin binding | Up-regulated in control vs treated (Everman et al. 2021) |
| *Jon99Ci* | proteolysis | endopeptidase activity, serine-type endopeptidase activity | ND |
| *Lectin 24Db* | ND | fucose binding, glycosylated region protein binding, mannose binding | Downregulated in control vs treated (Everman et al. 2021) |
| *NtR* | chemical synaptic transmission, ion transmembrane transport, nervous system process, regulation of membrane potential, signal transduction | extracellular ligand-gated ion channel activity, neurotransmitter receptor activity, transmembrane signaling receptor activity | ND |

**Figure 2.**
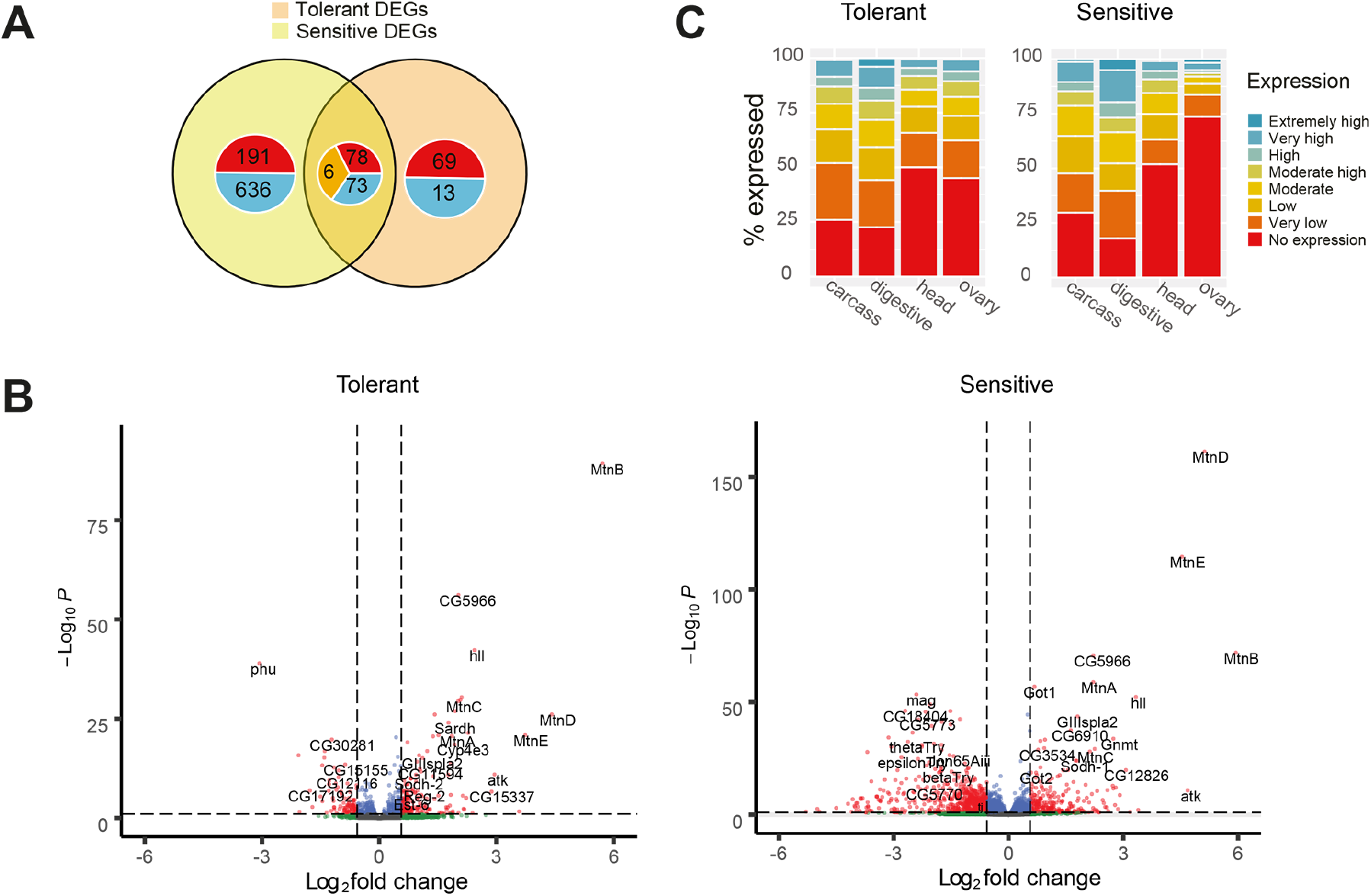
Copper differential gene expression and tissue analysis. **A)** Venn diagrams showing the degree of overlap between the differentially expressed genes in response to copper across the three tolerant and three sensitive strains. The numbers represented in red are found up-regulated, those in blue down-regulated and those in orange are the genes with discordant changes in expression between tolerant and sensitive strains (Table 1). Expression data obtained from mated female whole-body RNA-seq (3 replicates of 20 females each for treated and control conditions). **B)** Volcano plots of gene expression in tolerant (left) and sensitive strains (right). The horizontal dashed line represents the minimum adjusted p-value threshold (0.05), while the vertical dashed lines represent the fold change thresholds (log_2_(1.5) = 0.58). **C)** Classification of the tolerant DEGs and sensitive DEGs according to the levels of expression that these genes have in carcass, digestive system, head and ovary according to the DGET expression database.

As expected for metal treatment, the metallothioneins *MtnA-MtnE* were the most significantly differentially expressed genes by a large margin, both in tolerant and sensitive strains (Fig. 2B). While there was no relationship between their degree of induction and tolerance, all six strains were found to carry the 3’ indel polymorphism in *MtnA* that had previously been linked to oxidative stress resistance [17]. Other genes previously documented to play a role in copper homeostasis were notably absent from the differential expression lists, including the *Ctr1* family of transporters, *ATP7, Ccs* and *Malvolio*, suggesting that increased tolerance goes beyond metal chelation and homeostasis. Note that tolerant and sensitive strains did not differ in the expression of any of these genes in basal (nonstress) conditions either (Table S4B).

In order to find the tissues displaying the greatest levels of transcriptomic change after copper exposure in tolerant and sensitive strains, we used the *Drosophila Gene Expression Tool* (DGET, [38]). We classified our DEGs —taken from whole body samples— according to their degree of expression in four of the available DGET tissue databases: head, carcass, digestive system, and ovaries of four day old females (Fig. 2C). We focused on the overlap between our DEGs and those from DGET found to have higher levels of expression in these tissues (those categorized as having either high or extremely high expression: with RPKM values greater than 100). We found that the greatest level of overlap between DEGs and highly expressed genes according to DGET was seen for transcripts from the digestive system (hypergeometric test: tolerant p-value= 2.55×10^−19^; sensitive p-value = 9.59×10^−36^). The genes from our analysis that were highly expressed in the gut included the five metallothioneins *MtnA-E* and multiple serine peptidases, where many more peptidases were found significantly more down-regulated in sensitive than in tolerant strains. DEGs were also enriched to a lesser degree in the carcass (hypergeometric test: tolerant p-value= 4.58×10^−7^; sensitive p-value = 7.44×10^−14^). No DEG enrichment was seen for either the head or ovaries. Regarding gut subsections, the most notable overlap between our DEGs and highly expressed genes according to DGET were found in the copper cell region and the posterior gut (Fig. S2B). Copper cells are responsible for copper storage [39, 40], and changes in gut acidity [41]. One such marker of gut acidity —Vacuolar-type H+ATPase (*Vha100-4)* [42] *—* was found down-regulated by 0.6 (p-value = 0.01) and 2.0 (p-value =1.66×10^−8^) across tolerant and sensitive strains respectively, suggesting that gut acidity may be playing an important physiological role.

### Metabolism, Reproduction and Peptidase Inhibition Contribute to Copper Response in Tolerant and Sensitive Strains to Different Degrees

To determine what biological and physiological processes might differ between the tolerant and sensitive strains after the same period of copper exposure, we performed gene ontology (GO) enrichment analysis on tolerant and sensitive DEGs (Table S5A). Metabolism related terms were commonly seen as the largest and most significantly overrepresented terms in both tolerant and sensitive strains, although the exact processes often varied between the two groups (Fig. 3A). Chitin metabolic process (GO:0006030, adjusted p-value <0.0001 both in tolerant and sensitive strains) and Chitin binding (GO:0008061, adjusted p-value <0.0001 both in tolerant and sensitive strains) were also common to both analyses (Table S5A). As expected, response to metal ion (GO:0010038, adjusted p-value = 8.34×10^−4^) was strongly overrepresented in the copper tolerant strains (Fig. 3A). Additionally, several GO terms linked to reproduction and vitellogenesis—including vitelline membrane formation involved in chorion-containing eggshell formation (GO:0007305) and loss of vitellogen (encompassed by GO:0030704)— were found to be overrepresented in both analyses, but more so in sensitive strains (Fig. 3A and Table S5A). Note that shutdown of egg production is often a consequence of heavy metal toxicity [43, 44]. The majority of these terms were also found to be overrepresented in Gene Set Enrichment Analysis (GSEA) (Table S6). Further KEGG analyses also emphasized the role of protein metabolism processes in copper stress response and lysosome activity both in tolerant and sensitive strains (Table S7). Overall, these results suggest that under our assay conditions, the more tolerant strains could be undergoing metabolic stress response after 24 hours of copper treatment, while the more sensitive strains could be progressing to shutting down non-essential biological processes—such as egg laying—at the same stage, as has been previously described for other stress responses [45, 46].

**Figure 3.**
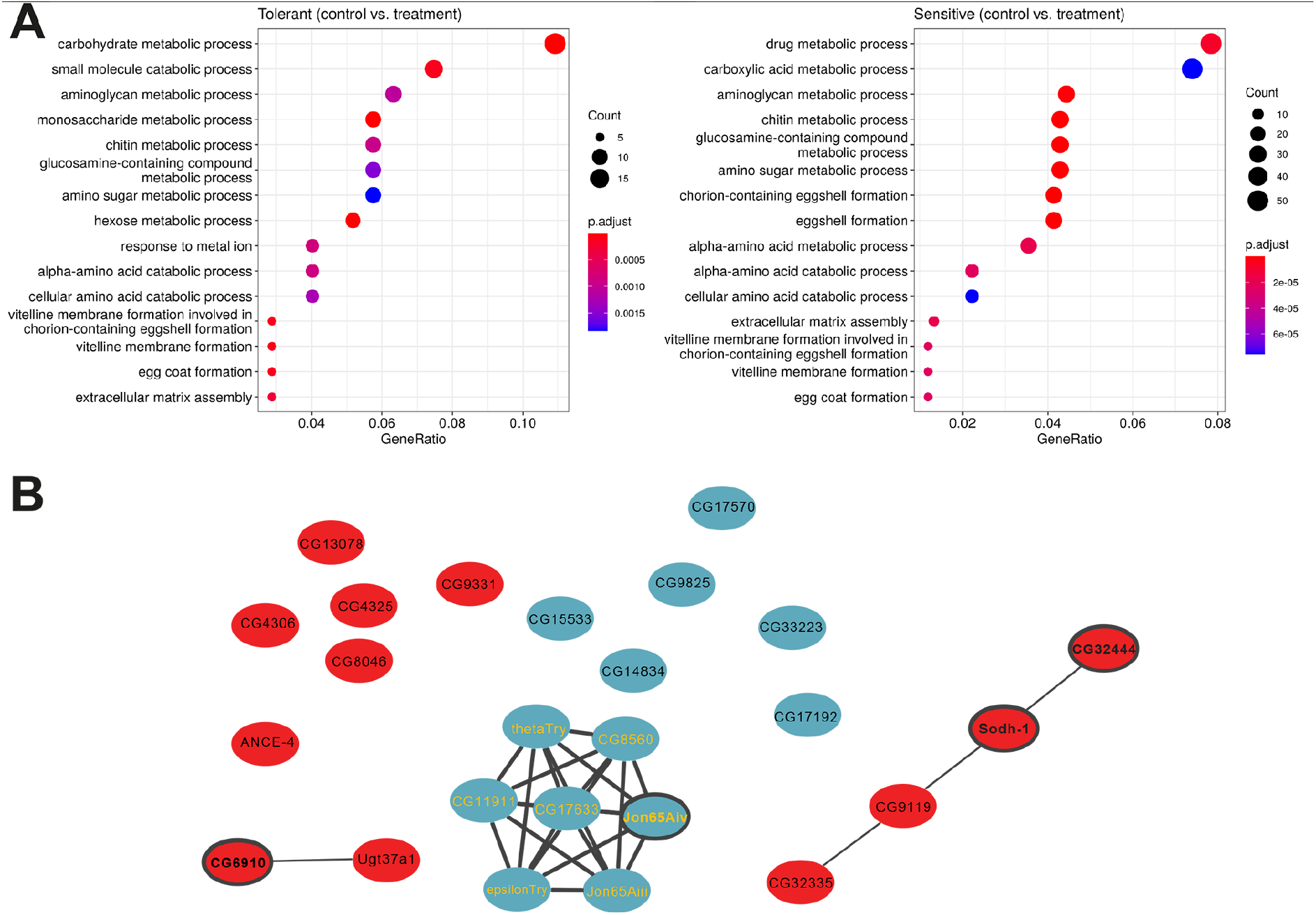
GO enrichment and correlational clustering analysis. **A)** Top 15 enriched GO terms associated with the DEGs in tolerant and sensitive strains. The Y-axis indicates gene functions, and the X-axis indicates the percentage of total DEGs in a given GO category (gene ratio). **B)** String interaction network of candidate genes taken from highly correlated modules (modules 2 to 5, Fig. S4) of the MCC analysis for the treated samples of the tolerant strains (PPI enrichment p-value: < 1.×10^−16^). Up-regulated genes are shown in red, down-regulated in blue. The genes with yellow labels are part of the serine peptidase cluster, while those with borders in bold were selected for further validation using RNAi knockdown or gene disruption lines.

Tolerant and sensitive strains also differed in basal gene expression (Table S4B), with the most significantly enriched molecular functions being enzyme inhibitor activity and endopeptidase inhibitor activity (Fig S3 and Table S5B). Thus, similar to other stress responses, differences in basal gene expression between tolerant and sensitive strains contribute to differences in copper stress responses between these strains (*e*.*g*. [47]).

Finally, we also investigated the level of gene co-regulation in response to copper of tolerant and sensitive strains using modulated modularity clustering (MMC) analysis, which in contrast to previous analyses does not rely on any prior gene functional annotations [48]. Tolerant strains show a high level of expression coordination after copper exposure while sensitive strains showed the opposite pattern (Fig. S4 and Table S8). Briefly, across the tolerant strains, we identified 24 modules with an average positive correlation, |r|, of 0.72 in treated samples, and 17 modules with a |r|=0.65 in controls. The higher correlations values and greater degree of partitioning observed in the treated samples indicated that there are coordinated changes happening after copper exposure (Fig. S4). For sensitive strains, 21 modules were identified in the treated samples, with an average |r|=0.77; and 40 in the controls with and |r|=0.71, with a less pronounced degree of partitioning, indicating a less coordinated response after 24 hours of copper exposure (Fig. S4). Heatmaps of the tolerant treated strains suggested that genes in modules 2-5 were very closely linked (Fig. S4, Table S8). Analysis of the 25 genes represented by these modules in STRING [49] revealed a group of seven tightly interacting serine peptidases (Fig. 3B) that are found highly expressed in the digestive system. While a number of these genes were from the *Jonah* family of serine peptidases, the discordantly expressed gene in tolerant vs sensitive strains, *Jon99Ci*, was not included among them (Table 1). Of these seven serine proteases, four have previously been shown to be regulated by the histone and protein deacetylase *Sirtin 2* (*Sir2;* [50]). On further inspection, 58 candidate DEGs from our tolerant strains and 187 from our sensitive strains were previously shown to display differential expression after *Sir2* knockdown, a significant overlap (hypergeometric test: tolerant p-value = 1.30×10^−20^; sensitive p-value = 6.87×10^−45^; Fig. S5A) [50]. Along with its role in heterochromatin formation, *Sir 2* is thought to have many additional protein targets that alter gene regulation. Among the targets of *Sir2*, gene expression datasets are available for *DHR96, dfoxo* and *HNF4* knockouts [51–53]. While a small degree of overlap was seen between our differential expression lists and that of *dfoxo* (hypergeometric test: tolerant p-value = 3.10×10^−7^; sensitive p-value = 1.50×10^−20^) and *DHR96* knock out analyses (hypergeometric test: tolerant p-value = 3.04×10^−7^; sensitive p-value = 3.58×10^−12^; Fig. S5), the greatest overlap was seen with *HNF4* knock out analyses (hypergeometric test: tolerant p-value = 1.45×10^−19^; sensitive p-value = 8.94×10^−66^; Fig. S5B) [51–53]. While little is known about the precise role *HNF4* plays in the midgut, its inferred link to the serine peptidases suggests a potential role in gut function.

### Eight out of 10 Copper Candidate Genes Were Confirmed to Play a Role in Copper Tolerance

Ten of the candidate genes associated with copper response based on our transcriptomic analysis were chosen for further characterization. Three of these genes —*CG5966, CG5773*, and *Cyp4e3*— were chosen on the basis of their differential expression data alone. Three other candidates—*Sodh-1, CG6910*, and *CG32444*—have been linked to copper homeostasis previously in the literature, but their exact functions have not been well characterized [12, 54, 55]. The remaining four candidate genes, *CG11594, Cyp6w1, Cyp6a8*, and *Jon65Aiv* were all found to be associated with TE insertions (see below). In addition, four of the ten candidates were part of the MMC cluster containing the serine peptidases (*Sodh-1, CG6910, Jon65Aiv*, and *CG32444*; Fig. 3B).

Seven of the genes tested showed changes in phenotype (copper survival) when knocked-down or disrupted in the direction that could be expected based on our RNA-seq data, *i*.*e*. if the gene was found to be up-regulated in response to copper, the knock-down of the gene was associated with decreased survival (Fig. 4, Fig S6, Table 2, Table S9). Mortality curves were significantly different for six of these seven genes when comparing the gene disruption or knock-down lines with their genetic background controls (Fig. S6 and Table 2), with four of them also showing significant differences in LT_100_ (Fig. 4 and Table 2). On the other hand, *CG6910* only showed differences in LT_100_ (Fig. 4 and Table 2). Of these seven confirmed genes, *CG11594, Cyp6w1, Cyp6a8*, and *Jon65Aiv*, are novel candidates, whose full role in copper biology is not yet understood, while the other three genes, *Sodh-1, CG6910*, and *CG32444* have prior links to the phenotype [12, 20, 55]. On the other hand, *CG5966* displayed decreased mortality when knocked-down, which was not predicted by its induction on copper (Fig. 4, Fig. S6, Table 2).

**Table 2.** Eight out of 10 Copper Candidate Genes Play a Role in Copper Tolerance. NS: not significant; ND: no data. Average mortality: RNAi, GDP (gene disruption) and Del (gene deletion): stock/background (Figure 4). ⬆: increased mortality. ⬇ decreased mortality. –: No data. Crosses with different backgrounds are separated with “;”. Reciprocal crosses results are separated with “,”.

| Gene | FC in expression after copper treatment |  | Kaplan-Meier |  |  | LT<sub>100</sub> |  |  |
| --- | --- | --- | --- | --- | --- | --- | --- | --- |
|  | Tol. | Sen. | RNAi | GDP | Del | RNAi | GDP | Del |
| *CG11594* | 2.06 | 1.84 | ↑<sup><0.001, <0.001</sup> | — | ↑<sup><0.001</sup> | ↑<sup>0.04, 0.004</sup> | — | ↑<sup>0.002</sup> |
| *Cyp6w1* | 2.03 | NA | ↑<sup><0.001, 0.028</sup> | — | — | ↑<sup>0.032, 0.059</sup> | — | — |
| *Cyp6a8* | NA | 1.91 | — | ↑<sup><0.001</sup> | — | — | ↑<sup><0.001</sup> | — |
| *Sodh-1* | 4.22 | 3.46 | ↑<sup><0.001</sup> | — | — | ↑<sup><0.001</sup> | — | — |
| *CG6910* | 2.86 | 3.09 | NS; NS | NS | — | NS; NS | ↑<sup>0.031</sup> | — |
| *Jon65Aiv* | -1.83 | -4.02 | ↓<sup>0.0062, <0.001</sup>,<br>NS | — | — | NS; NS | — | — |
| *CG32444* | 3.84 | 2.70 | ↑<sup><0.001</sup> | — | — | — | NS | — |
| *CG5966* | 4.07 | 4.64 | ↓<sup><0.001, <0.001</sup> | — | — | NS | — | — |
| *CG5773* | -2.24 | -5.13 | NS | — | — | NS | — | — |
| *Cyp4e3* | 3.81 | 2.05 | NS; NS | — | — | NS; NS | — | — |

**Figure 4.**
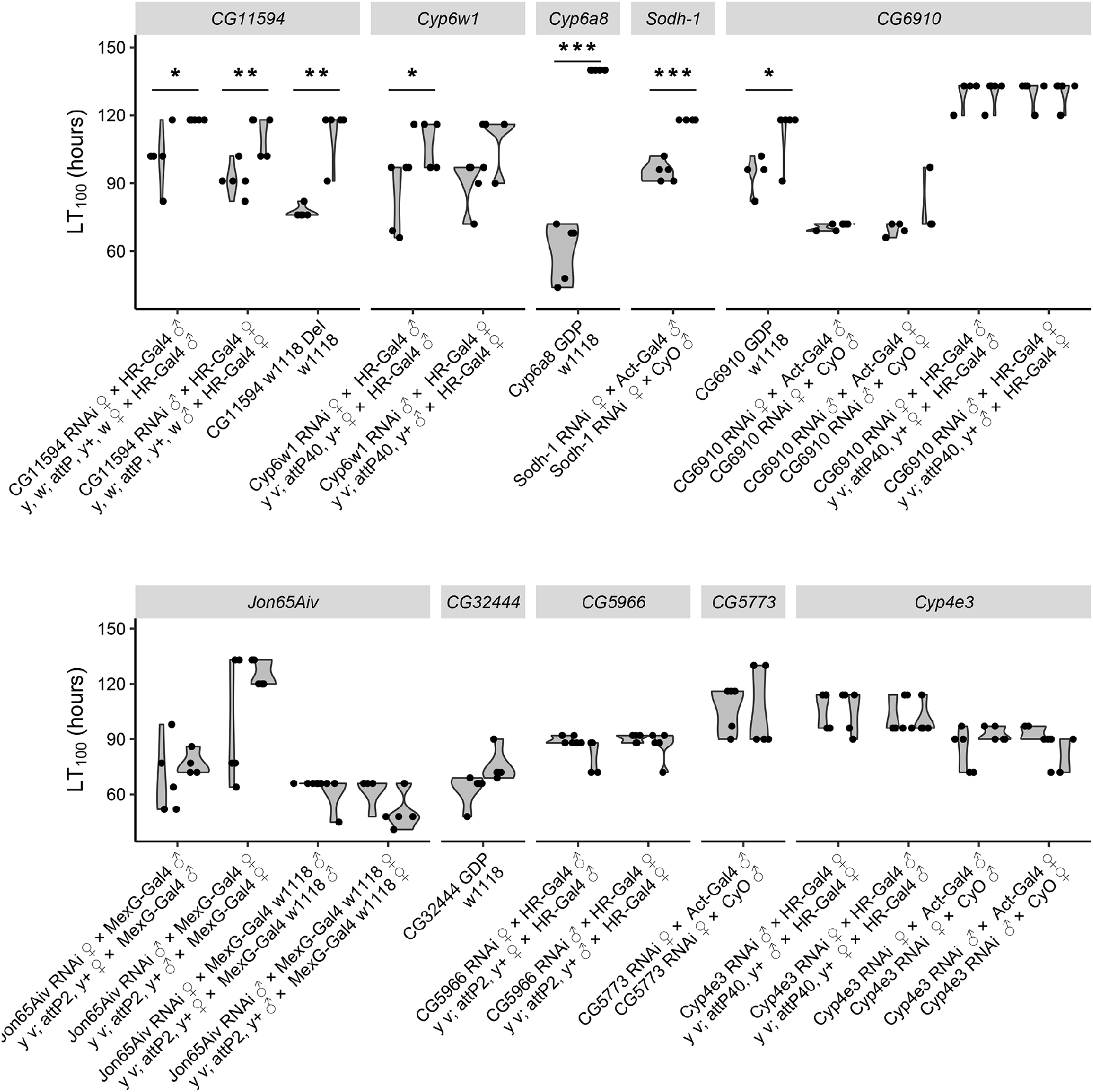
Copper survival experiments for all ten copper candidate genes. LT_100_ values comparing candidate gene disruption and knock-down lines with their genetic background controls (3 to 5 replicates of 15 females). Significant LT_100_ with p-values <0.05 are shown with *, p-values < 0.01 with **, and p-values < 0.001 with ***. Del is deletion and GDP is gene disruption. For RNAi knockdowns, genes thought to act in the ‘detox’ tissues—including the gut fat body and Malphigian tubules—were targeted with the *6g1HR-Gal4* driver, while those genes whose expression was more gut specific were targeted with *MexG-Gal4*. An ubiquitous *Actin5C-Gal4* knockdown was used for all other crosses. Drivers abbreviations are as follows: MexG-*Gal4(I) w1118* is the introgressed *w1118* version of MexG-Gal4 driver; *HR* is the *HikoneR* driver, and *Act* is the *Act5C* driver (see Table S1C).

*CG5773* and *Cyp4e3* did not display any changes in survivorship after knock-down (Fig. 4 and Table S9, Table 2). As *Cyp4e3* was initially tested with *6g1HR-Gal4* driver, based on prior expression data [56], we repeated the crosses with the *Actin5C-Gal4* driver. However, these additional assays did not show any significant changes in copper survival. While it is possible that the effects of these genes on copper tolerance are background sensitive, as per the example of *Cyp6g1* [57] and *Cyp12d1* [58], it is also possible that these genes have little to no true impact on the phenotype at all, and are only present due to co-regulation with other genes that do directly affect copper tolerance—a phenomenon that has been observed with regards to the *Cnc*/*Keap1* pathway [59] (Fig. 4, Fig. S6, Table S9).

### Copper Tolerance is Correlated with Gut Acidity and *CG11594* Activity and not Mitigated by Changes in Feeding Behavior

Both our DGET analysis and our GO analysis have lent evidence to the idea that there is a relationship between the gut and copper tolerance (Fig. 2C and Fig. 3A). As copper accumulation in *D. melanogaster* has previously been linked to changes in gut physiology [13], we assayed the changes in gut pH after copper exposure. Adults from the six RNA-sequenced strains were subject to copper assay conditions, and then allowed to recover for two hours on regular media supplemented with a mixture of Bromophenol Blue and yeast. If the acidic copper cell region of the gut remains un-inhibited by copper, this region should remain yellow under Bromophenol Blue (pH<2.3). After two hours, most recovering individuals had consumed enough media for the dye to be detected in the gut. Only AKA-018 had more than 10% of flies failing to feed on recovery– a phenomenon that was not seen in the controls (Fig. 5A). While all six strains showed decreased acidity across the copper cell region after copper treatment, the three sensitive strains showed a much higher loss of acidity than the three tolerant (Fig. 5A and Table S10A). Greater than 25% of individuals across all tolerant strains maintained a low pH under treated conditions compared to less than 10% of the sensitive strains (χ^2^ p-value = 4.815×10^−7^, Fig. 5A). Differences were less pronounced under control conditions, with only JUT-008 showing an appreciable loss of acidity in the absence of copper. From these results, we can infer a link between the loss of acidity in the copper cells and a decreased ability to tolerate copper.

**Figure 5.**
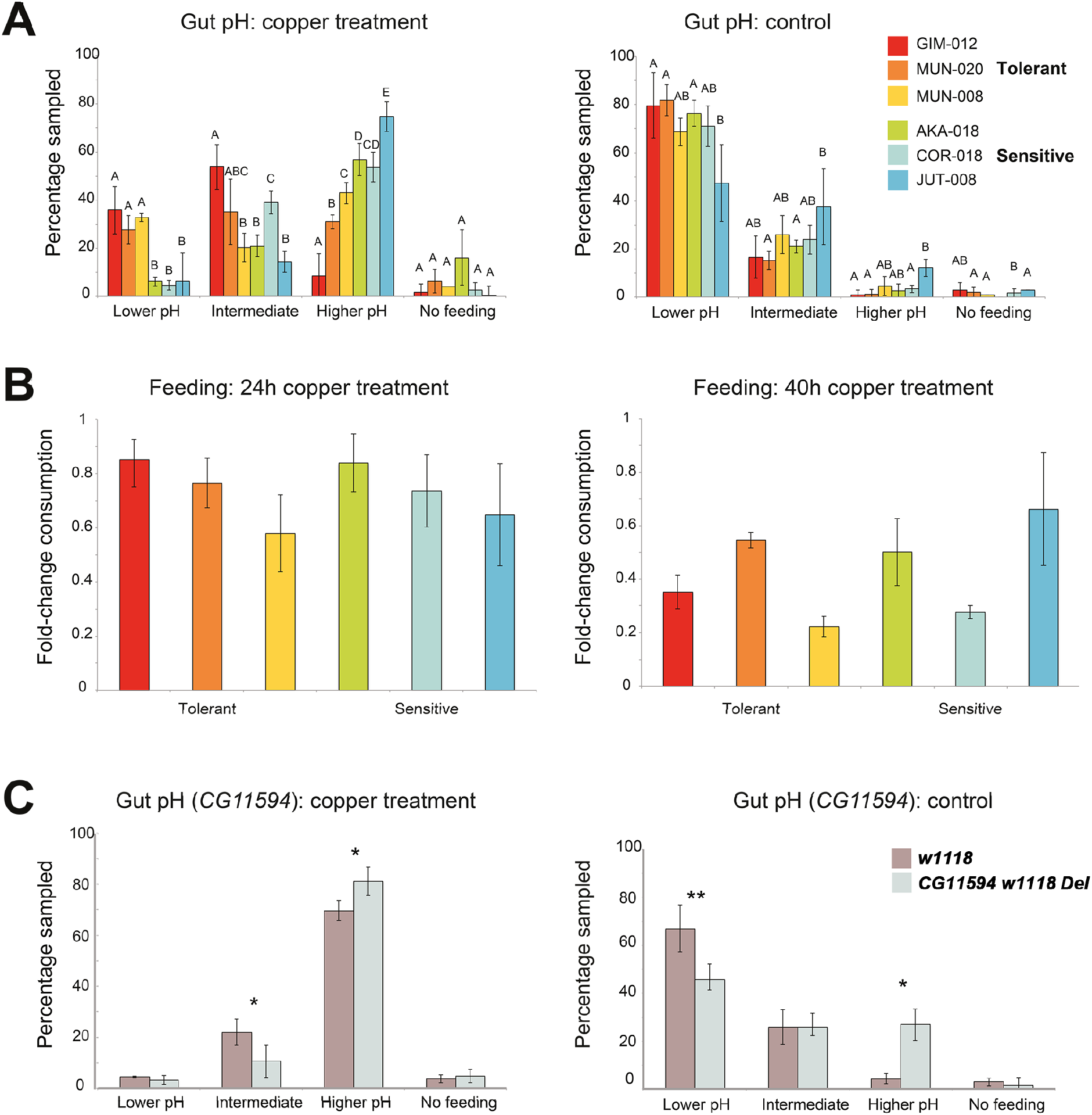
Gut acidity after copper exposure is correlated with copper tolerance and is not linked to feeding behaviour. **A)** Gut acidity results on the six RNA-sequenced strains after 24 hours of copper treatment and 2 hours of recovery (left) and after 24 hours of control conditions and 2 hours of recovery (right). Lower pH indicates that the dye turned yellow within the region of the gut containing copper cells; intermediate pH indicates that the dye turned green-brown, but a discrete acidic region could still be detected; higher pH indicates that the entire midgut was blue and the copper cell region could not be detected; and no feeding (clear or pale blue). The labels above the error bars (A-D) indicate significance across multiple tests - if a letter is shared between two bars in a grouping (*e*.*g*., Lower pH; Intermediate pH), the differences between said bars are not significant. **B)** Feeding avoidance in the presence of copper measured as a fold difference in consumption between treatment and control at 24 hours (left) and 40 hours (right). **C)** Gut pH of the control strain *w[1118]* (in red) and the *CG11594* deletion strain (in blue) after 16 hours of copper treatment (left) and control conditions (right). An asterisk (*) indicate a difference across the treatment groups at p-value < 0.05 (*) and p-value < 0.01 (**). For all the plots, error bars represent the standard error of the mean of three replicates containing 28-44 females each (A and C) and three replicates containing 25-30 females each (B).

*D. melanogaster* often avoid food sources with high concentrations of heavy metals [60, 61]. To determine if the changes in gut acidity are influenced by changes in feeding behavior, we repeated the copper tolerance assays on the six RNA-sequenced strains, this time with the addition of a 1% Erioglaucine Disodium Salt to both the treatment and control solutions to act as a dye. We measured the level of dye consumed in both treatment and control conditions at two separate time-points to determine the level of feeding avoidance. At 24 hours, the level of feeding avoidance on copper across most of the lines was quite low when compared to their control counterparts, with no significant differences between strains (Fig. 5B and Table S10B). Feeding avoidance was generally stronger at the 40-hour mark, where both MUN-008 and COR-018 were distinguishable by their greater levels of feeding avoidance (Fig. 5B and Table S10B). However, no relationship could be drawn between feeding behavior and whether or not the line showed high or low copper tolerance or changes in gut acidity.

If the observed changes in gut acidity are not based in behavior, they are likely physiological in nature. While metallothioneins could be good candidates [14], as mentioned above we found *MtnA-MtnE* to be up-regulated in response to copper both in tolerant and sensitive strains (Fig. 2B), and no differences in *MtnA-MtnE* expression between tolerant and sensitive strains were found in basal conditions (Table S4B). Thus, overall, changes in *MtnA-MtnE* expression are not likely to explain the identified differences in gut acidity (Fig. 5A). We thus decided to focus on *CG11594* one of the seven candidate genes that we confirmed as having a role in copper tolerance (Fig. 4 and Table 2), as this is the most poorly characterized of the candidate genes, which although being associated with a number of stress phenotypes it had no prior links to copper biology before this work [62, 63]. To determine if *CG11594* expression alters gut acidity, similar exposure and gut staining assays were carried out over a sixteen hour time period on a *CG11594* deletion strain (*w[1118]; CG11594[1]*), using *w[1118]* as the background control strain. While both lines displayed a high degree of gut de-acidification after treatment than any of the six natural lines, the effects seen on the *CG11594* deletion line were significantly greater than those on the background control line (p-value < 0.05, Fig. 5C and Table S10C).

Curiously, the clearest differences between the two lines were seen not in the copper treatment, but in the control, where only a half of the *CG11594* deletion individuals displayed a clearly defined acidic region. This is in stark comparison to the six sequenced strains, which displayed healthy guts under control conditions. These results suggest that physiology, not behavior, is the main driver behind midgut de-acidification after copper exposure, and that *GG11594* plays a role in this change.

### Transposable Element Insertions May Influence Copper Tolerance

TE insertions are often associated with changes in gene expression under stressful conditions (*e*.*g*. [64]), and in *D. melanogaster*, several specific insertions have been linked to stress response including zinc stress (*e*.*g*. [27–32]). However, until recently only the subset of TEs annotated in the reference genome could be analyzed, thus limiting the power of genome-wide analysis to investigate this type of structural variant. We took advantage of the availability of *de novo* whole genome assemblies and *de novo* TE annotations for the three tolerant and three sensitive strains analyzed in this work [24], to investigate the association between proximal *cis* TE insertions and gene expression levels in both treated and control conditions (within 1kb of the insertion, see Material and Methods). Using QTLtools [65], we identified three TE insertions that were significantly associated with changes of expression in nearby genes: two in response to copper (*FBti0061509* and *FBti0063217*) and one both in control and in response to copper (*FBti0060314*; Table S11A). Although the number of significant associations is small, this is most probably due to the small number of genomes analyzed (six)— suggesting that this approach should provide more insight with larger datasets.

As an alternative approach, we also investigated whether previously identified DEGs in tolerant and sensitive strains were located within 1kb of a TE insertion (Table S11B). There were no significant differences between the percentage of differentially expressed genes located within 1kb of a TE in tolerant compared to sensitive strains (14.28% across the three tolerant strains and 11.29% in the three sensitive; p-value= 0.2193). While 73.5% of the TE insertions were associated with gene up-regulation in tolerant strains only 28% of the TEs were associated with up-regulation in sensitive strains (Fisher’s exact test p-value = 0.0014; Table S11B). Because the effect of transposable elements, and other genetic variants, are often background dependent (*e*.*g*. [66]), we also investigate whether TEs were associated with DEGs identified at the strain level. None of the strains showed a significant enrichment of TEs nearby DEGs (test of proportions p-value > 0.05, Table S11C).

Finally, we tested three TE insertions for their effects on copper tolerance. For each of the TE insertions, we constructed two outbred populations: one with the insertion and one without the insertion (see Material and Methods). This strategy limited testing to those TE insertions that have been found segregating in populations at a high enough level that we could obtain enough strains to construct the outbred populations. We chose two insertions that besides being located nearby DEGs, showed signatures of positive selection in their flanking regions suggesting that they might be adaptive: *FBti0020036* and *FBti0020057* [26]. The third TE candidate, *FBti0020195*, is not present in any of our six sequenced strains but garnered special interest due to its location within *CG32444*, a candidate gene identified in this study and further confirmed with the use of gene disruption (Fig. S6). For each of these three TE insertions, we constructed two outbred populations: one with the insertion and one without the insertion (see Material and Methods). For each of the paired outbred populations, those containing TE insertions demonstrated greater survivorship on copper than their negative counterparts, both on LT_100_ (p-value = 0.01117, p-value < 0.001 and p-value = 0.0018 for *FBti0020036, FBti0020057* and *FBti0020195* respectively, Fig. 6 and Table S11D), and across the entire survival curve (log-rank tests p-value < 0.001 for all three comparisons, Fig S7 and Table S11D).

**Figure 6.**
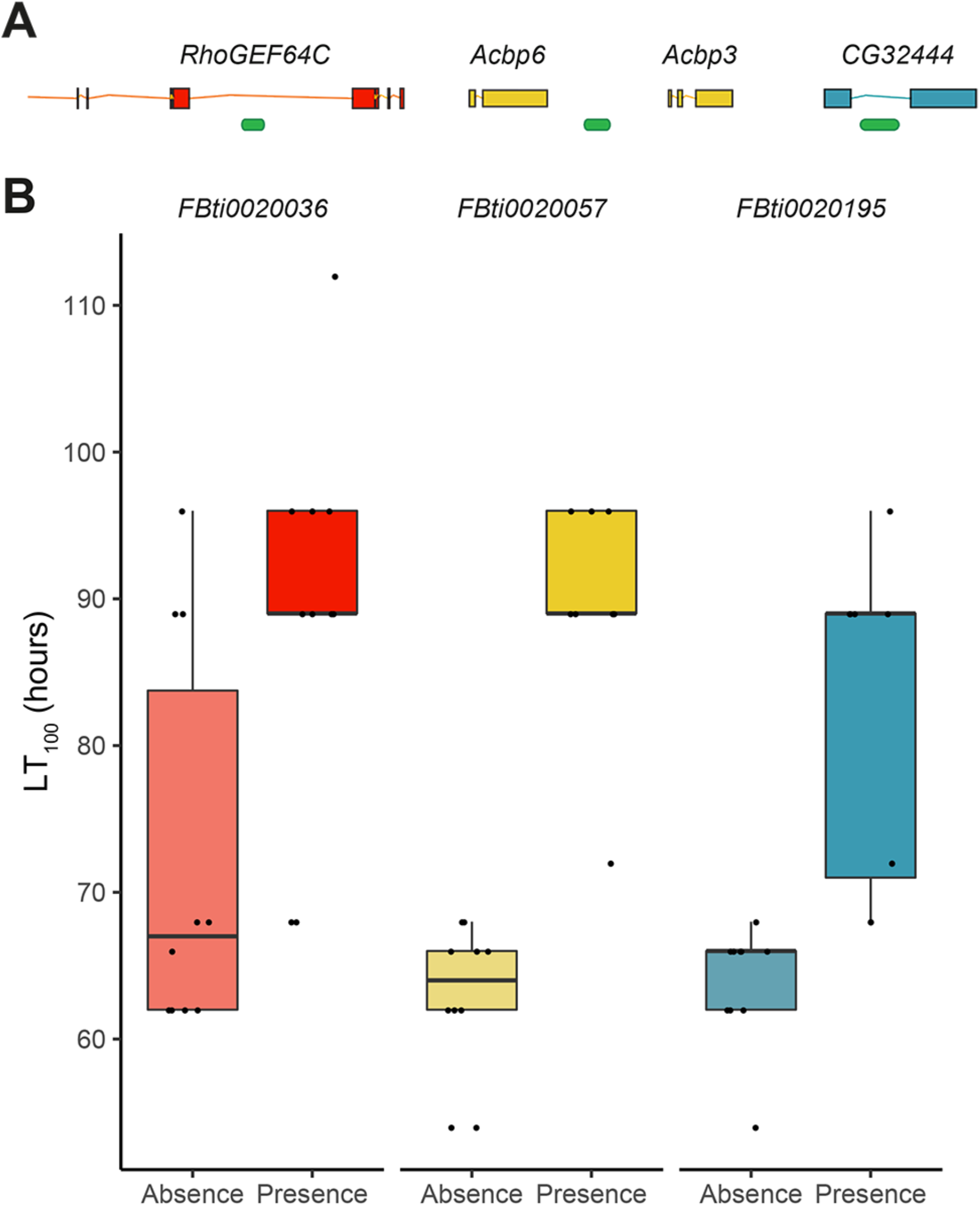
Copper survival experiments for the three candidate transposable element insertions. **A)** Gene structure showing where each of the candidate TEs are inserted. For *RhoGEF64C* only the 3’ region of the gene is depicted. **B)** Relative change in average mortality at the end of the assay comparing outbred populations with and without the candidate TE (9 to 10 replicates of 10-15 females in treatment and 5-10 females in control conditions).

## DISCUSSION

### The Environmental Determinants of Copper Tolerance in *D. melanogaster* are Complex

In this study, we undertook a survey of multiple European *D. melanogaster* populations to determine how copper tolerance varies across the continent, and whether this variation could be linked to the presence of copper or other environmental factors. To achieve this, we compared our phenotypic values with geographic factors, copper soil levels, atmospheric pollution levels, and degree of urbanization. We found a positive correlation between latitude and LT_50_ (p-value = 0.015, Table S3B). While we also found evidence of a link between urban build-up and greater tolerance, no clear relationship could be drawn between tolerance and any of the direct measures of pollution available to us. As Romic and Romic (2003) [2] noted, human sources of environmental copper are characterized by many point sources of contamination, and while we are aware that some well-known sources — such as atmospheric copper— are missing from our dataset, it is possible that there are others missing as well. Moreover, it is also unknown whether the greatest effect will be from an accumulation of multiple sources of the metal, or a small number that are the most bio-available. As these point sources can be difficult to characterize, performing environmental sampling, *e*.*g*., soil sampling, alongside fly collections may be a viable alternative [67]. The diversity of vegetation may also be worthy of record as copper uptake and storage varies across plant tissues and species [68]. Although we cannot discard that more extensive sampling could further help discern the relationships between phenotype and environment, our results indicate that the finer details of the surrounding environment should be receiving as much attention as the finer details of the genome when making sense of phenotypic differences.

### The Genetic Basis to Copper Tolerance in *D. melanogaster* is Complex, and Involves Multiple Regulatory Factors

One of the most distinguishing features of our phenotypic dataset is the high degree of variation both within and between sampling locations (Fig. 1B and Fig. S1). While high levels of phenotypic variation can sometimes result from an allele of large effect segregating within a population, as seen in Battlay et al. (2016) and Green et al. (2019) [58, 69]; the gradual distribution of our LT_50_ values suggest that this is not the case and that the degree of phenotypic variation seen across our strains is likely an indication of the polygenic basis of the trait (Fig. S1A, [5, 20]). This was in turn backed by our RNA-sequencing analysis, which indicated that copper tolerance is a trait with a complex genetic architecture, involving multiple genes and regulatory factors, and with a large degree of expression change occurring in the gut.

With regards to genes with prior links to metal response, variation in metallothionein expression was not found linked to phenotypic variation in the six strains sequenced (Fig. 2B). However, as all six strains carry the 3’ indel that is believed to be linked to increased stress tolerance, and it is found to be close to fixation in northern Europe [17], it is likely that metallothionein tolerant variants have already been subject to selection. We also saw no significant differences in expression with regards to multiple genes previously linked to copper homeostasis. While this may initially come across as curious, many of the previous studies characterizing copper-related genes in genetically modified lines were carried out in a small number of strains with the aim of characterizing genes that play a role in human diseases [9, 55, 70], and not explicitly copper exposure in nature. While the lack of these genes in our DEG lists does not necessarily mean that they are not involved in copper tolerance, it does indicate that the genes contributing to the variation we see in tolerance in natural populations of *D. melanogaster* are much broader than previously characterized in these studies, and that the biological basis behind copper tolerance may be constrained by the need to maintain copper levels in less extreme environments.

While it has been well documented that MTF-1 plays an important role in regulating gene expression in response to metal exposure, including metallothionein induction, it is unlikely to be the only regulatory factor affecting changes in expression, especially with regards to downstream metabolic processes affected by copper toxicity [22]. By using a combination of DGET and gene clustering, we were able to identify *Sir2* and *HNF4* as additional potential regulatory elements. *Sir2* plays a multifaceted role in maintaining energy homeostasis, affecting fat mobilization [71], insulin signaling [72], and energy consumption [50]. *HNF4*—a direct target of *Sir2* regulation—also influences a wide range of processes involved in cellular metabolism and systemic physiology [53]. These results are supported by our functional gene analysis. Of the eight confirmed candidate genes (Fig. 4), *Sodh-1* and *CG32444* have both been linked to the kind of metabolic processes modulated by *HNF4* and *Sir2*, while also having found associated with copper toxicity previously [20, 55]. The two cytochrome P450s, *Cyp6w1* and *Cyp6a8*, are both linked to oxidative stress [58, 73], a process that has been linked to metal tolerance previously [22]. *CG6910* is down-regulated in MTF-1 knockout mutants [12]. The roles of the remaining three candidates in copper tolerance are more speculative. *Jon65Aiv* is known to be a serine protease with a likely role in digestion [74]. Serine proteases have also been shown to be down-regulated in clusters after exposure to another metal, manganese [75] and during aging [76], although the reason for this perturbation remains unresolved. As copper inhibits larval midgut acidification [14], a phenotype also seen in aging [13], it would be tempting to investigate the relationship between acidity and serine proteases directly. This also has interesting implications for cross species comparisons: While serine protease function is well conserved across species [77], the degree of segmentation and the pH levels of the alimentary tracts of many other insect species (*e*.*g*. lepidoptera) are not [78, 79].

While its role has not been well characterized, *CG5966* is involved in triglyceride breakdown [80], and starvation response [81], a functional profile that fits with regulation by both *Sir2* and *HNF4. CG5966* has also been found to be highly up-regulated during mitochondria dysfunction [82], along with many other stress response genes. Finally, our pH assays give the greatest guidance to the role of *CG11594*, which may prove to play a role in gut integrity.

While our individual gene candidates may not be so well conserved outside of *Drosophila, Sir2* and *HNF4* do have well-conserved orthologs, much in the same manner as the metallothioneins. While there is no previous evidence for these genes playing a role in copper toxicity in arthropods, such evidence exists in mammalian cell culture: rat hepatocytes treated with copper sulfate display increased expression of *Sir2* homologs *Sirt1* and *Sirt2* [83], while *HNF4-α*, influences copper responsive transcription changes in HepG2 cells [84]. Furthermore, while many of our punitive candidates for *Sir2 and HNF4* regulation were found highly expressed in the gut, both regulatory elements have been shown to play different roles in different tissues [53], presenting us with the possibility that not only might their roles in copper response be discordant in different tissues, but that this may apply to the general transcriptional signature post metal exposure as well. Future assays using knockdown or disruption of these factors across multiple tissues in *Drosophila* would be able to confirm their specific roles in copper response.

Further changes in gene expression can potentially be traced back to transposable element insertions. TE insertions are often associated with the differential expression of nearby genes under stress conditions [27, 28, 64]. We identify several TE insertions located inside or nearby differentially expressed genes (Table S10B). For three of these insertions we further showed that their presence is associated with increased copper survival (Fig. 6). Further analysis, such as recombination mapping and CRISPR based knock-outs in these genetic backgrounds could potentially assist in confirming the role of these specific TE insertions in altering gene expression and their effect on phenotype.

### Gut acidity is linked to copper tolerance in *D. melanogaster*

Our analysis demonstrated that a large degree of the differential expression observed after copper exposure was occurring in the gut, a key tissue when it comes to copper physiology [13–15]. A role for the gut is also supported by the GO enrichment results: chitin binding and metabolic processes suggest a role for the peritrophic membrane [85], which is important for gut integrity. A study on the effects of Lufenuron—a chitin disrupter— in *Anthonomus grandis* showed that gut disruption could lead to changes in metabolism and the down regulation of vitellogen; also seen in our GO enrichment analysis [86]. In addition, chitin binding and metabolic processes also affect the cuticle, which may affect copper exposure via contact. Indeed, copper DEGs were also found to be enriched amongst extremely high and highly expressed genes from the carcass in DGET (Fig. 2C). A correlation between cuticle darkening and increased body copper content has also been reported in *D. melanogaster* [75].

Our gut pH assays clearly demonstrate that copper exposure results in a loss of acidity in the copper cell region —and that this effect is more sharply seen in the three sensitive strains (Fig. 5A). Our subsequent feeding response assays excluded differences in copper consumption as a potential explanation of varying losses in gut acidity, suggesting a more physiological process was responsible for the changes observed (Fig 5B). While metallothioneins could be good candidates [14], our *Mtn* expression data do not sufficiently explain the differences we observed (Fig. 2B and Table S4). This opens up the possibility that one or more of our gene candidates selected for further analysis may be affecting copper tolerance through changes in copper cells or gut acidity. While the function of *CG11594* has mostly gone uncharacterized, its expression has been linked to both oxidative stress and ER stress in the DGRP [62, 63]. While disruption of *CG11594* expression caused a strong loss in gut acidity after copper treatment, there was a notable loss under control conditions as well (Fig. 5C). These results imply that loss of gut acidity is a sub-phenotype to copper tolerance, and that both share links to *CG11594* activity —although the exact mechanism underpinning the relationship remains elusive. In light of previous studies, we can propose two tentative alternative hypotheses: regulation of *CG11594* by both *Sir2* and *HNF4*, suggests that the gene plays a general role in energy and metabolism [50], and it is differences in the allocation of energy and resources that affects survival. Alternatively, links to ER stress [63] could indicate a role linked to lysosome function or metal storage.

## CONCLUSIONS

Our investigation across European natural populations of *D. melanogaster* proved copper tolerance to be a highly variable trait. We confirmed the involvement of multiple new candidate genes, identified two potential new regulatory factors that have previously only been seen to mediate metal responses in mammals, and described physiological changes linked to this trait. Unlike previous candidates, such as the metallothioneins, which are common across a wide phylogeny, it is unlikely that the exact genes shown to affect copper tolerance in *D. melanogaster* will be perturbed in other species vulnerable to metal toxicity. However, other, more general, molecular pathways and physiological changes in the gut we observed in *D. melanogaster* are likely to prove relevant in studying the effects of copper toxicity in other species.

## MATERIAL AND METHODS

### Fly Collections

Details of all the stocks used can be found in Table S1. The nine original collections were carried out across the summer of 2015 by the DrosEU consortium. Each of the established isofemale strains (4 to 16 depending on the population, Table S1) was repeatedly inbred for up to 20 generations. Of the additional strains included for geographical and environmental analysis, the Mauternbach and Recarei strains were caught in 2018 and the Bari strains were caught in 2011 and have been kept as isofemale strains since then [87]. All fly collection sites are documented in Fig. 1A. All strains were maintained on semolina-yeast-agar media and were kept at 25 °C on a 12:12 hour light and dark cycle for at least one generation before use.

### Copper tolerance assays

Copper sulfate (CuSO_4_) (CAT# 451657-10G) was obtained from Sigma Aldrich. Copper assays were adapted from Bonilla-Ramirez *et al*. (2011) [61]. This particular method was chosen for two reasons: (i) as *Drosophila* are known to show food avoidance with a high concentration of heavy metals [61] boib it allowed exposure via both contact and digestion; and (ii) the 4-5 day length of the assay gives sufficient time to differentiate between tolerant and sensitive strains without risking high control mortality.

Briefly, powdered CuSO_4_ was reconstituted to 20mM in a 5% sucrose solution. Brilliant blue food dye (E133) was added to aid visibility and even dispersal. An identical control solution without CuSO_4_ was prepared in the same manner. 250µl of the CuSO4 sucrose solution was pipetted onto 70×17mm slips of filter paper (Whatman, CAT# 3030917), which were then placed into 15ml Falcon tubes (Cultek, CAT# 352096), containing 1ml on 1% agar at the bottom. Papers were allowed to dry for 15 minutes before the flies were added. To assist respiration, holes were made in the lids of the falcon tubes. Number of dead flies was counted at different timepoints both in the control and treated conditions, until all flies were dead in the treated conditions.

For each isofemale strain, 4-7 day old females were used in the copper survival assays both in control and treated (20mM copper) conditions. Three replicates of up to 15 flies each were performed for the treatment and for the control conditions (Table S1). LT_50_ calculations were used to interpolate measures of survival for each of the strains. Linear models were fitted to time point-mortality data on a log-probit scale using the “glm” function in the R statistical package, using a script adapted from Johnson *et al*. (2013) [88]. Of the 73 DrosEU strains screened, LT_50_ values were successfully calculated for 71, along with the 26 additional strains from Italy, Austria and Portugal (Table S1).

### Correlation Analysis with Geographical and Environmental Variables

Copper soil concentration data was taken from The European Soil Data Centre (ESDAC: https://esdac.jrc.ec.europa.eu/content/copper-distribution-topsoils) [3], with the exception of the Tenerife data, which was taken from Fernandez-Falcon *et al*. (1994) [89]. Air pollution data was taken from the European Environment Agency (EEA): https://discomap.eea.europa.eu/map/fme/AirQualityExport.htm. The pollutants considered included: PM10 (particulate matter 10 micrometers or less in diameter), PM2.5 (particulate matter 2.5 micrometers or less in diameter); arsenic in PM10, cadmium in PM10 and lead in PM10 data. All measures were taken from the closest research station available for each catch site. General PM10 and PM2.5 data and atmospheric metal data for arsenic, cadmium and lead were available for the majority of catch sites (Table S3). Data for particulate copper taken from PM10 measures had to be excluded due to both insufficient geographical coverage and a lack of consistency in the measures made (PM10 and precipitation). All tests and linear regression models were performed in *R v3*.*5*.*1*. [90]. Regression models were fitted with LT_50_ values as the dependent variable, and with geographical and pollution measures as independent variables. Degree of urbanization of the fly collection locations was based on whether the closest population to a collection site was a city (> 50,000 inhabitants: urban), a town with a population > 5,000 inhabitants (semi-urban), or less dense populations <5,000 inhabitants (rural; Table S3). This degree of urbanization is based on the OECD/European Commission (2020), Cities in the World: A New Perspective on Urbanisation, OECD Urban Studies, OECD Publishing, Paris, available at: https://www.oecd.org/publications/cities-in-the-world-d0efcbda-en.htm) [37]. We performed a multiple linear regression model to test the association between copper tolerance (LT_50_) and the geographical and environmental variables. We first created a linear model with all the measured variables (model: LT50 ∼ Longitude + Latitude + Copper + PM10 + PM2.5 + Arsenic + Cadmium + Lead + DegreeUrbanization). We carried out a backward stepwise regression to eliminate variables using the *dropterm()* function of the MASS package in R. At each step we removed the least significant variable. Only variables with a p-value < 0.1 were retained in the minimal model [91], which considered Latitude and Degree of urbanization (R2=12%, p-value = 0.0079).

### RNA-seq Sample Preparation

RNA-seq analysis for short-term copper exposure (24 hours) was performed on six inbred strains, where those with the strain codes GIM-012, MUN-020 and MUN-008 of were copper tolerant and JUT-008, COR-018 and AKA-018 were copper sensitive (Appendix 1 and Table S1). To maximize odds of choosing mostly homozygous strains, we prioritized those strains with a high degree of inbreeding (minimum of F20), and a low degree of variation between biological replicates in the LT_50_ assays.

Four biological replicates of 25 mated female flies 4 to 7 day-old from each line—separated 24 hours beforehand on CO2— were exposed to CuSO_4_ or the equivalent control conditions, as reported above, and removed after 24 hours. This timeframe allowed low levels of death in the sensitive strains, but enough time to stress tolerant strains, as measured by the induction of *MtnB* detected through RT-qPCR. Deceased individuals from strains COR-018 and JUT-008 were removed before whole-body RNA extraction. 20 females from each biological replicate were flash frozen in liquid nitrogen and total RNA was isolated using the GenElute Mammalian Genomic RNA miniprep kit (Sigma Aldrich, CAT# RTN350-1KT), following the manufacturer’s instructions. For each sample, the three repeats with the best RNA quality based on BioAnalyzer were retained for sequencing. 1 µg of total RNA from each sample (whole female body) was used for subsequent library preparation and sequencing using an Illumina Hiseq 2500. Libraries were prepared using the Truseq stranded mRNA library prep according to the manufacturer protocol. Only two control samples for both AKA-018 and MUN-020 showed high enough quality for further sequencing analysis. Thus overall, we used 34 samples.

### Analysis of RNA-seq Data

RNA-seq analysis was performed using the *rnaseq* pipeline (v1.2) from the *nf-core* community, a *nextflow* collection of curated bioinformatic pipelines [92, 93]. The total number of raw reads obtained per sample range between 25,16 M and 46,13M. Briefly, sequencing quality was assessed using *FastQC* (*v*.*0*.*11*.*8*, [94]). *TrimGalore* (*v*.*0*.*5*.*0*) was used for adapter removal [95], and *Cutadapt v. 1*.*18* with default parameters was used for low-quality trimming [96]. Trimmed reads were mapped to the *D. melanogaster* genome r6.15 [97] using *STAR* (*v*.*2*.*6*, [98]). On average, 95.9% of the reads mapped to the reference genome. Technical duplications were explored using *dupRadar* [99]. Overall, we found no bias towards high number of duplicates at low read counts, so we did not remove duplicates from the alignments. We used *featureCounts* (*v*.*1*.*6*.*2*, [100]) for counting the number of reads mapping to genes (*reverse-stranded* parameter). Multi-mapping reads and reads overlapping with more than one feature were discarded. The matrix of counting data was then imported into *DESeq2* [101] for differential expression (DE) analysis following the standard workflow and applying the design formula: *Strain + Treatment* in the analysis of the tolerant and sensitive strains. To compare resistant vs. tolerant strains in basal conditions, we used the design formula ∼ *Resistance*. Normalization was performed using the standard DESeq2 normalization method, which accounts for sequencing depth and RNA composition [101, 102]. Differentially expressed genes were chosen based on both log_2_ fold change (> 1.5) and adjusted p-values (< 0.05). Gene counts and scripts to perform the DE analyses can be found at https://github.com/GonzalezLab/Dmelanogaster_Copper. Functional profile analyses of the differentially expressed genes (GO, GSEA and KEGG) were performed using the R package *clusterProfiler* [103]. Breakdown of differentially expressed genes by tissue was performed using the *Drosophila Gene Expression Tool* (DGET: https://www.flyrnai.org/tools/dget/web/; [38]), with gut subsampling data taken from similar aged flies from Marianes and Spradling (2013) [41].

Modulated Modularity Clustering (MMC) was used to group differentially expressed genes into subsets of genetically correlated genes in both treated and control samples. All analyses were carried out as outlined in Stone *et al*. (2009) [48], except the variance filtering, which was performed in *R v3*.*5*.*1* beforehand. The variance filter removed genes where no variance across repeats and samples was found, which basically removes genes with no expression. Additional network-based analysis was performed using *STRING* (v10, [49]) using a minimum interaction score of 0.7. Subsequent visualizations were performed using *Cytoscape* (v.3.7.1, [104]).

### RNAi and Gene Disruption Assays

Candidate genes were functionally validated using RNAi knockdown lines from the KK library of the *Vienna Drosophila Resource Centre* ([105]; obtained from the VDRC) and the Transgenic RNAi Project (TRiP) developed from the Harvard Medical School ([106]; obtained from the *Bloomington Drosophila Stock Centre*). Additional gene disruptions were performed using either *Drosophila Gene Disruption Project* (GDP) lines ([107]; obtained from the *Bloomington Drosophila Stock Centre*) and one independent deletion mutant ([w1118]; CG11594[1]; *Bloomington Drosophila Stock Centre)* All stock numbers are provided in Table S1.

The choice of *GAL4* driver was based on data obtained for each gene from *FlyAtlas 2* (http://flyatlas.gla.ac.uk/FlyAtlas2/index.html; [56]). Three of the drivers were homozygous: the *6g1HR-Gal4* driver, described by Chung *et al*. (2006) [108], and two different background versions of the *MexG-Gal4* driver, originally described by Phillips and Thomas (2006) [109]. All three were provided by Shane Denecke. The heterozygous *Actin:5C-Gal4/CyO* driver was obtained from Bloomington Stock Centre (BDSC ID 4144).

For all assays using homozygous GAL4 drivers, the mortality of all GAL4-RNAi crosses was compared to matching control crosses using the appropriate RNAi background strain. For the KK RNAi lines, comparisons were made to crosses using the KK construct-free control strain (VDSC ID 60100). For all assays containing TRiP RNAi lines, for those lines with the RNAi construct inserted into the attP2 site, comparisons were made to the y, v; attP2, y+ construct-free control strain (BDSC ID 36303) and or those lines with the RNAi construct inserted into the attP40 site, the y, v; attP40, y+ construct-free control strain (BDSC ID 36304). Due to difficulties maintaining the strain, for all assays using crosses using the *Actin:5C-Gal4/CyO* driver, the offspring that inherited the Gal4 construct were compared to their *CyO* inheriting siblings. All GDP lines and the [w1118]; CG11594[1] strain were compared to w1118.

Copper survival experiments were performed as described above using 4-7 day old flies. Three to five replicates of up to 15 flies for the treatment, and four to five replicates of up to 10 flies were performed for control conditions. Kaplan-Meier survival analysis was chosen as the best statistical comparison for comparing disrupted and control samples, and all analyses were performed using the R package Survminer (v.0.4.8). Additionally, relative change in average mortality is also provided as a proxy of the size of the effect of these genes on copper tolerance, significance was tested using a t-test.

### Gut pH Assays

4-5 day old flies from the six strains taken from the RNA-seq analysis were subject to the same assay conditions used for the copper tolerance assays for 24 hours. Assays were performed in triplicate, with each replicate consisting of 30-50 female individuals. Higher numbers were required for COR-018 and JUT-008 to account for the level of mortality expected during this timeframe. Flies were then transferred to regular *Drosophila* media, on which 200µl of a mixture of 1% Bromophenol Blue, dried yeast and water (at 1:1:3 ratio) had been added twenty minutes prior. Flies were permitted to feed for two hours before having their mid-guts dissected in PBS and accessed for loss of acidity. 28-44 samples were dissected from each replicate (numbers varied as guts were often very fragile). Any individuals who proceeded to die after transfer to recovery media were discarded. Samples were determined to have experienced minimal loss in acidity if the cells in the acidic region of the midgut remained yellow (pH < 2.3), an intermediate loss if they had faded to green or brown, and full loss if they could not be distinguished from the surrounding sections (pH > 4). No feeding was recorded if no media was present in the gut.

Similar assays were carried out on lines *w[1118]* and the *CG11594* deletion line, (*w[1118]; CG11594[1]*) (Table S1) over a shorter 16 hour time-period, to account for the greater sensitivity of these lines to copper.

### Feeding Avoidance Assays

To measure the effect that the presence of copper has on feeding avoidance, the six strains from the RNA-seq analysis were assayed in similar conditions to that of the copper tolerance assay, but with the addition of Erioglaucine Disodium Salt (1%, Sigma-Aldrich CAT#861146) to both the treated and control solutions. Eoglaucine Disodium Salt been shown to be an effective tracer up to 48 hours in *Drosophila* [110]. Assays were performed in triplicate for groups of 25-30 4-7 day old females, with higher numbers used for COR-018 and JUT-008 to account for the degree of mortality expected at the end of this time period. All dead individuals were discarded. Flies were homogenized using a pestle, with each sample consisting of three flies in 620µl of distilled water. After crushing, samples were spun at 14,000rpm for 10 minutes and then frozen for 24 hours. 180ul of supernatant was loaded into each well of a 96 well Nunc-Immuno™ MicroWell™ plate (Sigma-Aldrich, CAT#M9410). Measurements were made using a Techan Infinite® 200 Microplate Reader, at 630nm, after 10 seconds of agitation at 9mm. Three technical replicates for six samples, for a total of 18 wells, were loaded for each treatment condition. Each plate contained four water blanks and five standards containing between 0.015 and 1.5×10^−5^ % of dye. The amount of dye consumed was inferred from a linear model fitted from the points of the standard curve. All results are reported as the fold difference in feeding between treated and control samples for each time point.

### Transposable Element Analysis

#### eQTL analysis

The RNA-Seq data for tolerant and sensitive strains both in control and treated conditions were trimmed using the fastp package (v.0.20.0) [111] with default parameters. Expression levels were quantified with the salmon package (v.1.0.0) [112] against the ENSEMBL (Dm.BDGP6.22.9) transcripts. Obtained transcripts per million (TPM) were summed up to gene level and rlog normalized using DESeq2 (v.1.28.1) [101]. To test the association between gene expression and TE variants, we used the TE annotations for each one of the six genomes analyzed available at https://github.com/sradiouy. The genotype table with the information of the presence/absence of all the TEs present in each one of the strains was created using custom script (https://github.com/sradiouy).

The eQTL analysis was performed using the QTLTools package (v.1.2) [65]. Putative cis-eQTL for the six strains were searched within a 1 kb window around each gene using the cis module in QTLTools in control and in treated conditions separetely. No trans effects were considered. We used the nominal pass to evaluate the significance of the association of each gene expression level to all the TE insertions within the 1kb window. This nominal pass involves the testing of all possible variant-phenotype via linear regression. The variant-phenotype pair with the smallest nominal p-value is kept as the best QTL for that particular TE. In addition, we also performed a permutation pass (100,000 permutations) to adjust for multiple testing. We focused on the significant TE-gene associations with a with a nominal p-value < 0.05 and and an adjusted p-value < 0.05.

#### Identification of TEs Nearby DEGs

Reference gene annotation was lifted over to each of the six strain assemblies analyzed using Liftoff (v.1.4.2, [113]), with default parameters, to produce gene annotations of each strain in GFF format. Liftoff annotation was transformed to BED format with a custom python script (https://github.com/sradiouy). Then bedtools closest (v2.29.2, [114]) was used to define TE insertions within a 1kb of each gene (parameters: -k 10, -D ref) using the TE annotations available at https://github.com/sradiouy. We used the prop.test() function of R to assess whether there is an enrichment of TEs in DE genes compared to the whole genome for each strain.

#### Phenotypic Validation

TE present and TE absent outbred populations were constructed for three candidate insertions: *FBti0020195, FBti0020057* and *FBti0020036*. Each of these outbred populations was developed to have a mixed genetic background, while remaining consistently homogenous for either the presence or absence of the selected element [115]. For each outbred population, ten females and ten males from each of the five nominated strains (four in the case of *FBti0020195+*) were pooled to establish each population (Table S1). Each outbred was maintained for 8 generations in cages before being screened. Copper tolerance assays were carried out as per the prior experiments, using 4-7 day old females. 9 to 10 replicas of up to 15 flies in treated and up to 10 flies in control were performed for each outbred population (Table S10C). The experiment was run until all flies were dead. Kaplan-Meier survival analysis was performed on present and absence pairs in the same manner as above. Relative change in average mortality is also provided as a proxy of the size of the effect of these genes on copper tolerance.

## DECLARATIONS

### Ethics approval and consent to participate

Not applicable

### Consent for publication

Not applicable

### Availability of data and materials

Data is available in the supplementary tables. Genome assemblies and the raw data (long and short read sequencing) have been deposited in NCBI under the BioProject accession PRJNA55981. RNA-sequence data is available under NCBI accession number: PRJNA646768; GEO: GSE154608. The six sequenced genomes are available together with gene, TE annotations and RNA-seq coverage profiles generated in this work for visualization and retrieval through the DrosOmics genome browser [116].

### Author contributions

L.G. contributed to the design of the work, to the acquisition, analysis and interpretation of data and to the drafting the manuscript. M.C-Z., S. R. and G.E.R. contributed to data analysis and to the revision of the manuscript. J.S-O. contributed to the design of the work, to the acquisition of data and revised the manuscript. J.G. conceived the study, contributed to the design of the work, to the acquisition, analysis and interpretation of data and to the drafting the manuscript. All authors approved the submitted version of the manuscript.

## Acknowledgments

We thank DrosEU members for sharing the European strains, and Shane Denecke and Trent Perry for sharing GAL-4 driver lines. We thank Joshua Schmidt for scripts related to the Kaplan-Meier analysis. We thank Luciano Massetti for making us aware of the availability of the atmospheric pollution data.

