## Supplementary Figures for "The genomic basis of copper tolerance in *Drosophila* is shaped by a complex interplay of regulatory and environmental factors"

**Figure S1. Copper tolerance phenotypes across all populations.**

**A)** LT_50_ values for the 71 isofemale collected in 2015 by the DrosEU consortium (see Table S2A for the name of each strain). The majority of these strains were inbred between 15-20 generations before screening (Table S1A). **B)** LT_50_ values for the 26 isofemale strains from additional locations in Austria, Portugal and Italy collected in 2011 and 2018. The strains in each chart are arranged in order of ascending LT_50_ (see Table S2A). Bar colour corresponds to copper concentration, as per the map in Fig. 1A. The cooler colours refer to low copper concentrations and the warmer colours refer to high copper concentrations. Error bars represent 95% CI of the probit slope. The pink labels in **A)** represent the three sensitive strains and the blue labels represent the three tolerant strains that were subject to subsequent differential expression analysis and genome sequencing.

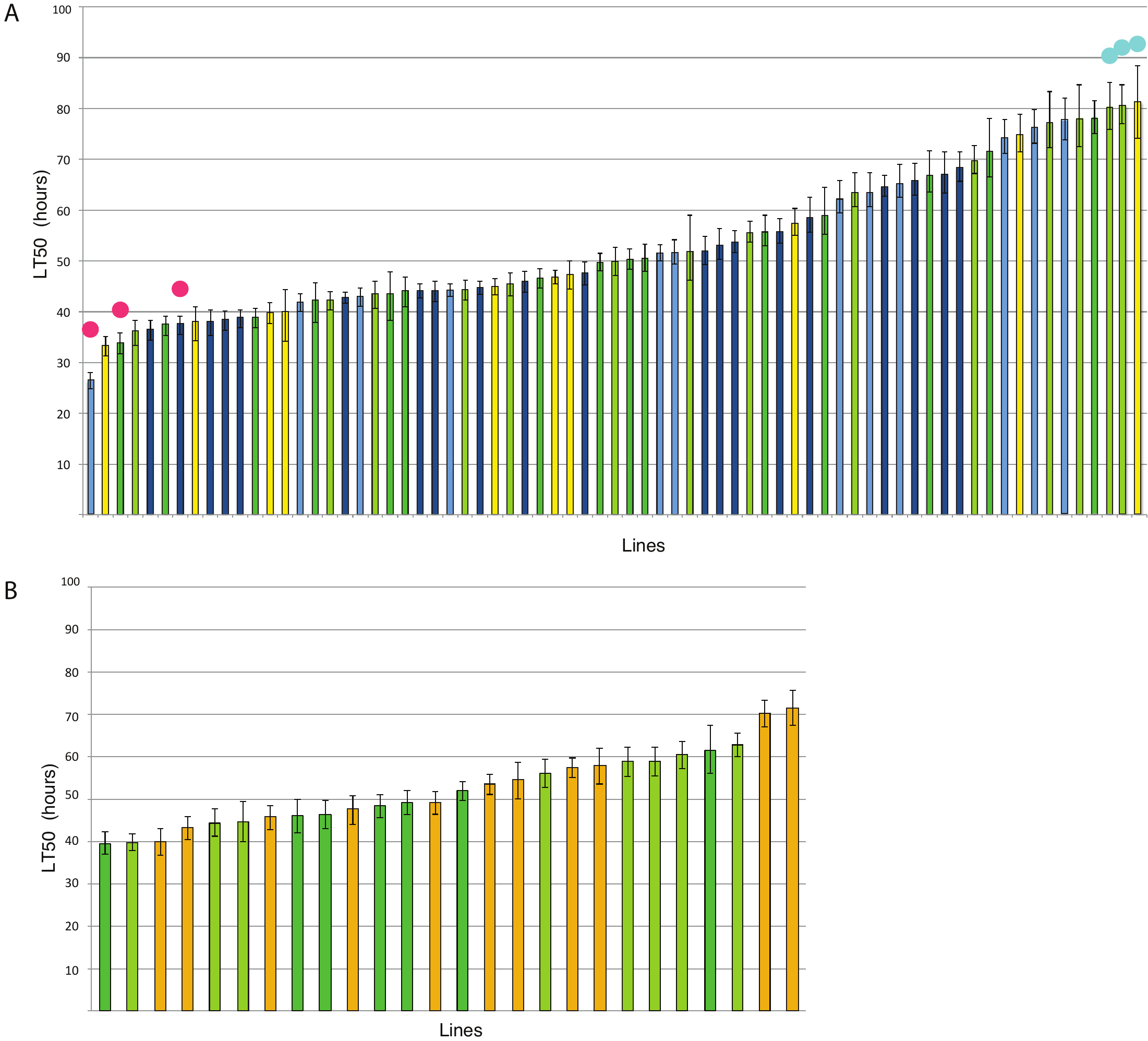

**Figure S2.**

**A)** Principal component analysis of expression data in both treated and control conditions for the six analyzed strains. Darker shades for each line, with a xxCx label represent control samples, lighter shaded points with xxDx labels represent copper treated samples. **B)** DGET expression analysis for gut subsections. Breakdown of gene expression levels for tolerant (top) and sensitive (bottom) strains. Subsections: a = Anterior (regions 1-3); Cu = Copper Cells; Fe = Iron Cells; LFCs = Large Flat Cells; p = Posterior (regions 1-4). With the exception of the a1 segment of the gut in tolerant strains, genes found highly and extremely highly expressed in all sub-sections were significantly enriched for our DEGs.

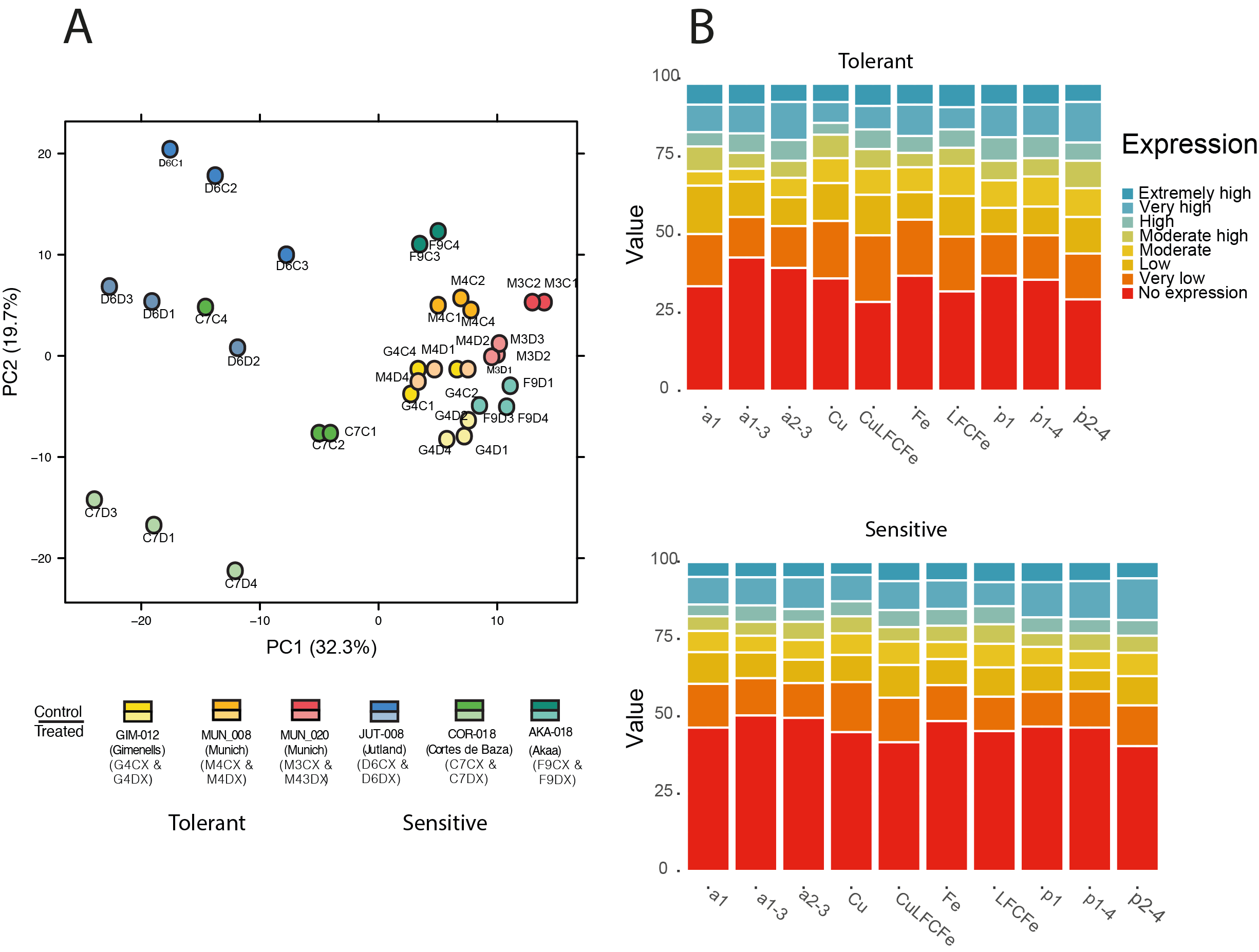

**Figure S3. GO clustering analysis for DEGs between tolerant and sensitive strains in basal conditions.**

Top enriched GO terms associated with the DEGs when comparing tolerant *vs.* sensitive strains under control conditions. The *y*-axis indicates gene functions, and the *x*-axis indicates the proportion of total DEGs in a given GO category (gene ratio).

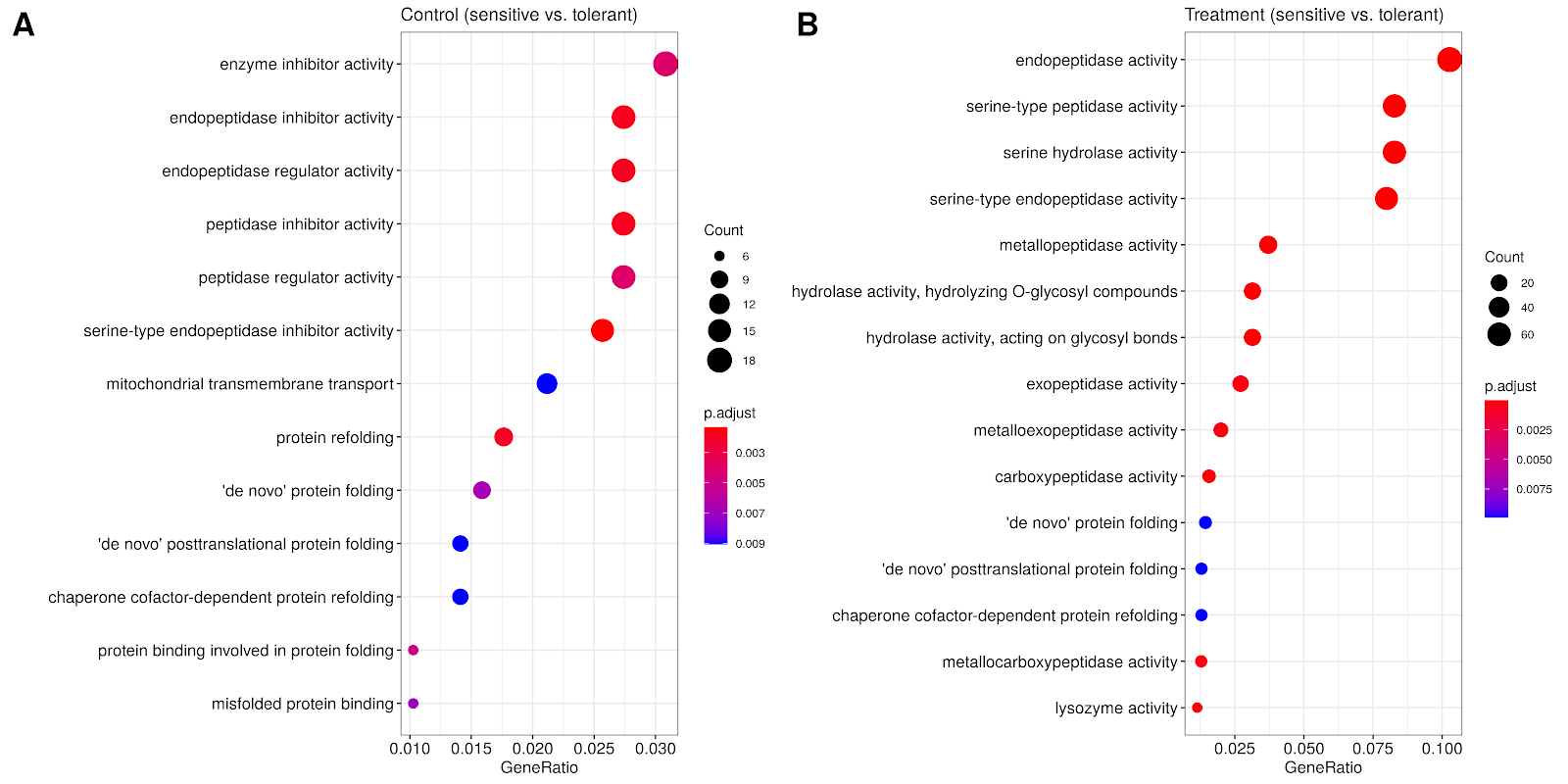

**Figure S4. MMC analysis of DEGs before and after copper exposure.**

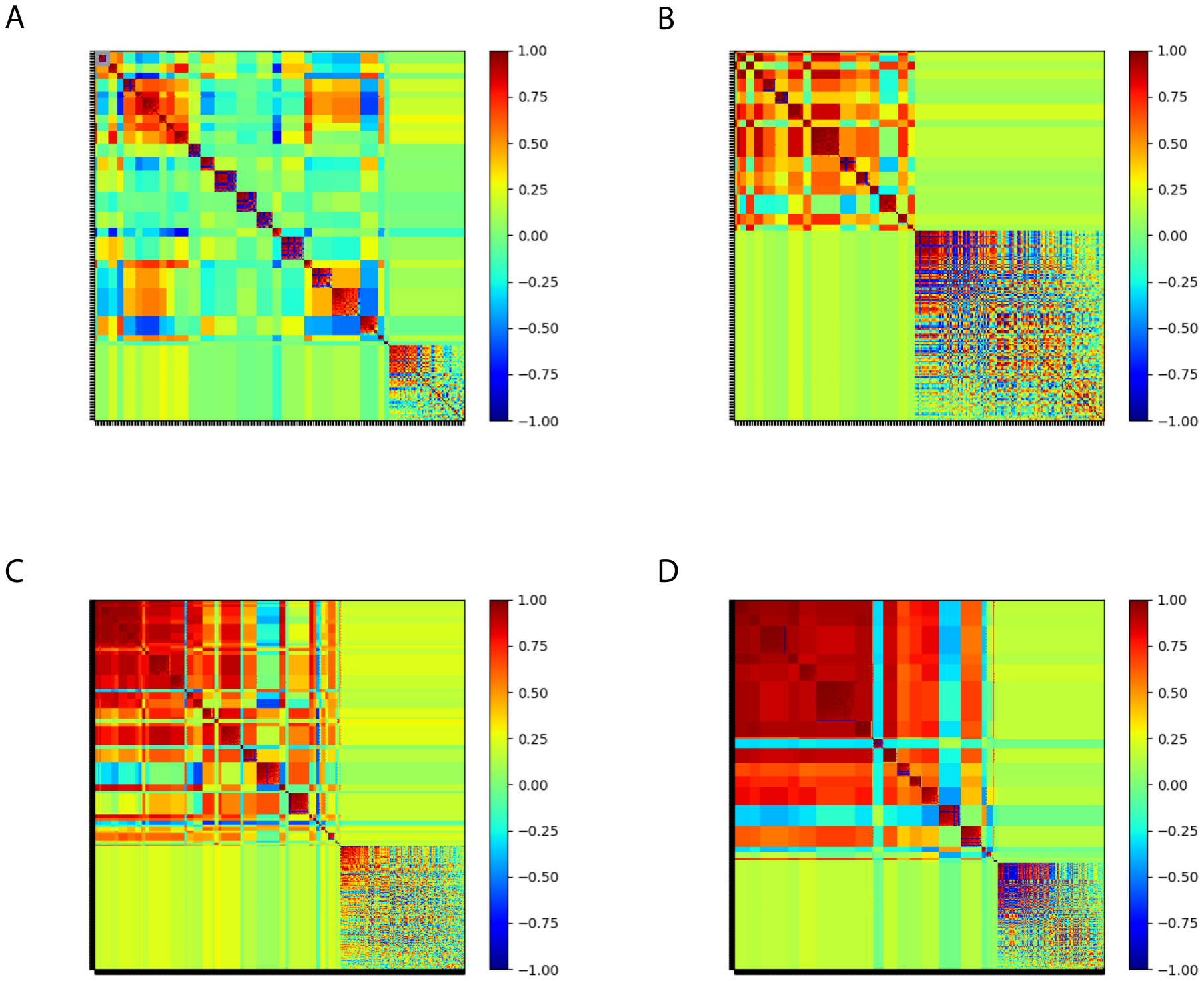
Analysis was performed separately for both tolerant (**top**) and sensitive (**bottom**) strains. Treated samples are shown on the left (**A, C**) and control on the right (**B, D**). Each coloured point represents the Spearman correlation (*r*s) between two genes.
Across the tolerant strains, we identified 24 modules with an average positive correlation, |r| , of 0.72 from treated samples, and 17 modules with a |r| = 0.65 from controls, indicating a higher level of expression co-ordination after copper exposure. For sensitive strains, 21 modules were identified in the treated samples, with an average |r| = 0.77; and 40 in the controls with and |r| = 0.71, with a less pronounced degree of partitioning, indicating that expression in sensitive strains is actually less modulated after 24 hours of copper exposure (see also Table S8).

**Figure S5. Overlapping expression patterns between copper DEGs and regulatory factors knockout and knockdowns.**

**A)** Genes responding to copper exposure overlap with those regulated by *sir2* and *HNF4.* Venn diagrams showing the degree of overlap between the DEGs between *Sir2* and *HNF4* knockouts along with tolerant (left) and sensitive strains (right). **B)** Venn diagrams showing the degree of overlap between the DEGs for two additional downstream targets of *sir2*. Overlap between DEGs from tolerant strains and *DHR96* knock-outs (top left), sensitive strains and *DHR96* knock-outs (top right), tolerant strains and *dFoxo* knock-outs (bottom left) and sensitive strains and *dFoxo* knock-outs (bottom right). The numbers represented in red are found commonly up-regulated, those in blue commonly down-regulated and those in yellow are discordant.

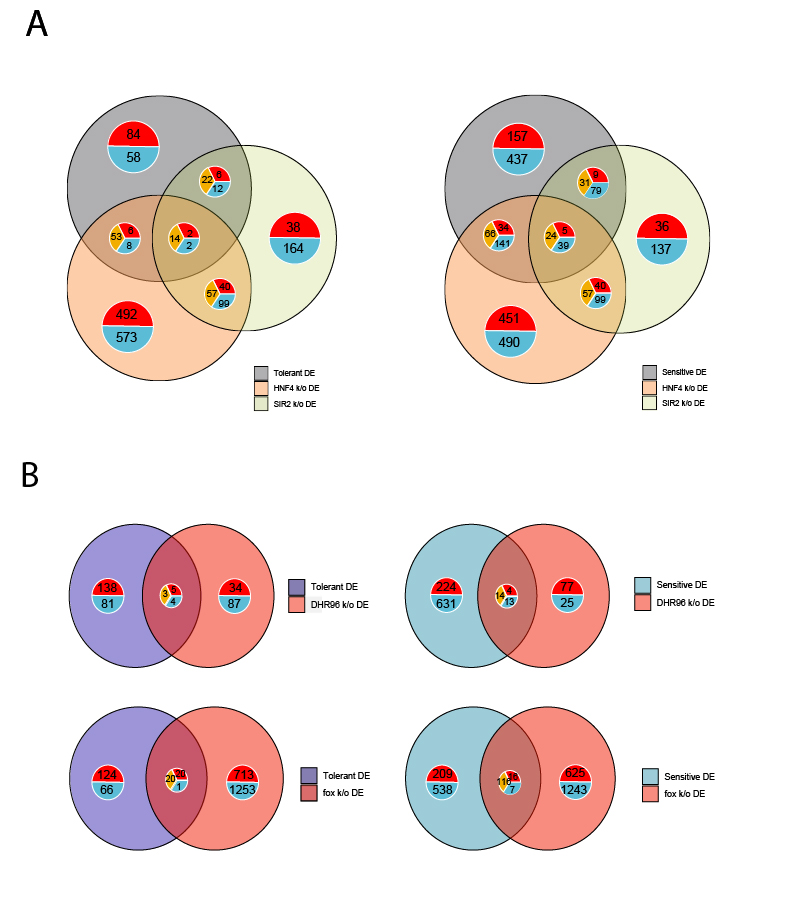

**Figure S6. Kaplan-Meier survival curves for the survival assays performed on RNAi knockdowns and disruption for all gene candidates.**

Shaded regions indicate the 95% confidence intervals. Statistical significance was estimated by using log-rank tests. Those plots shaded in blue are the knockdown or disruption lines and plots in red are the control lines.

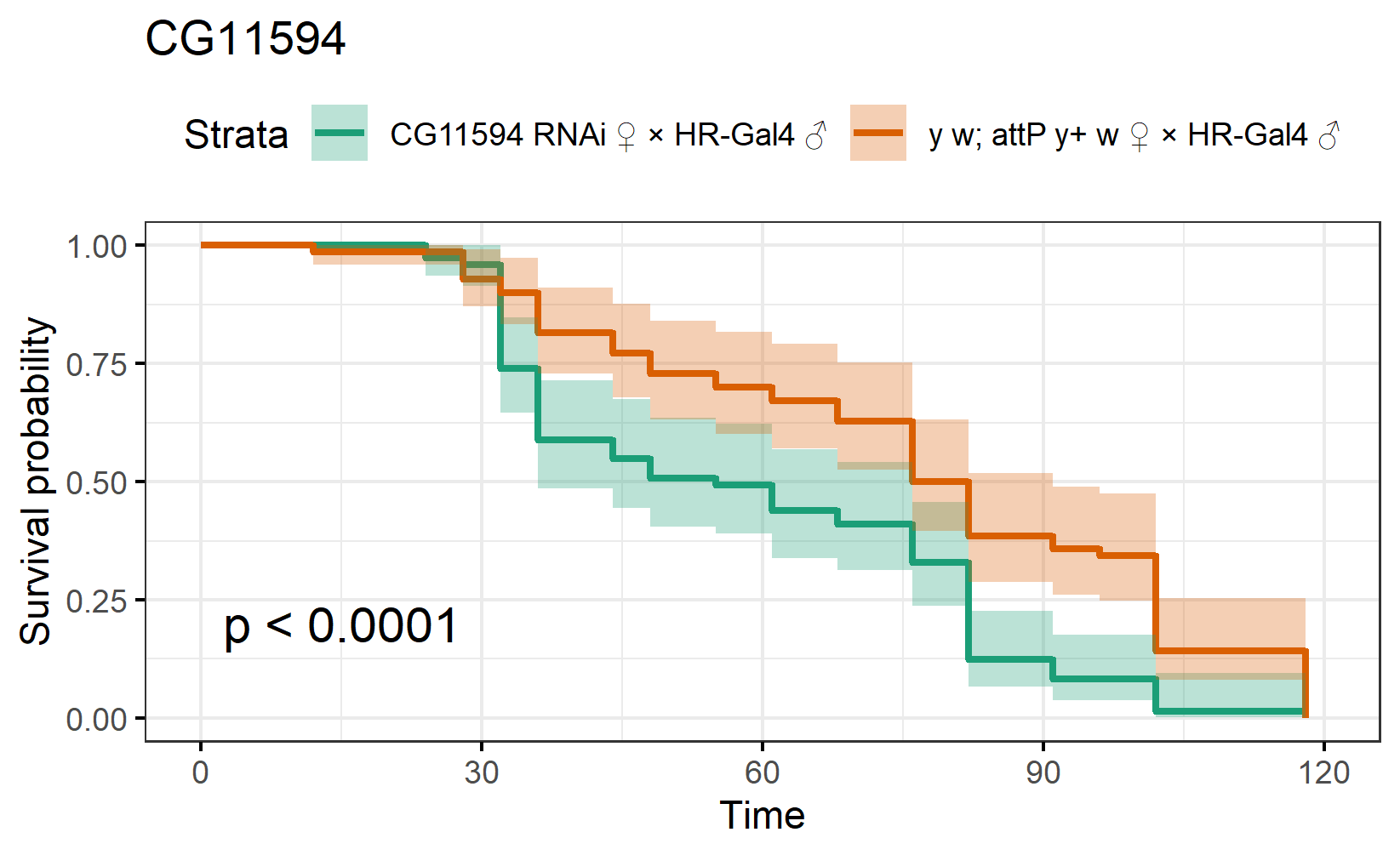

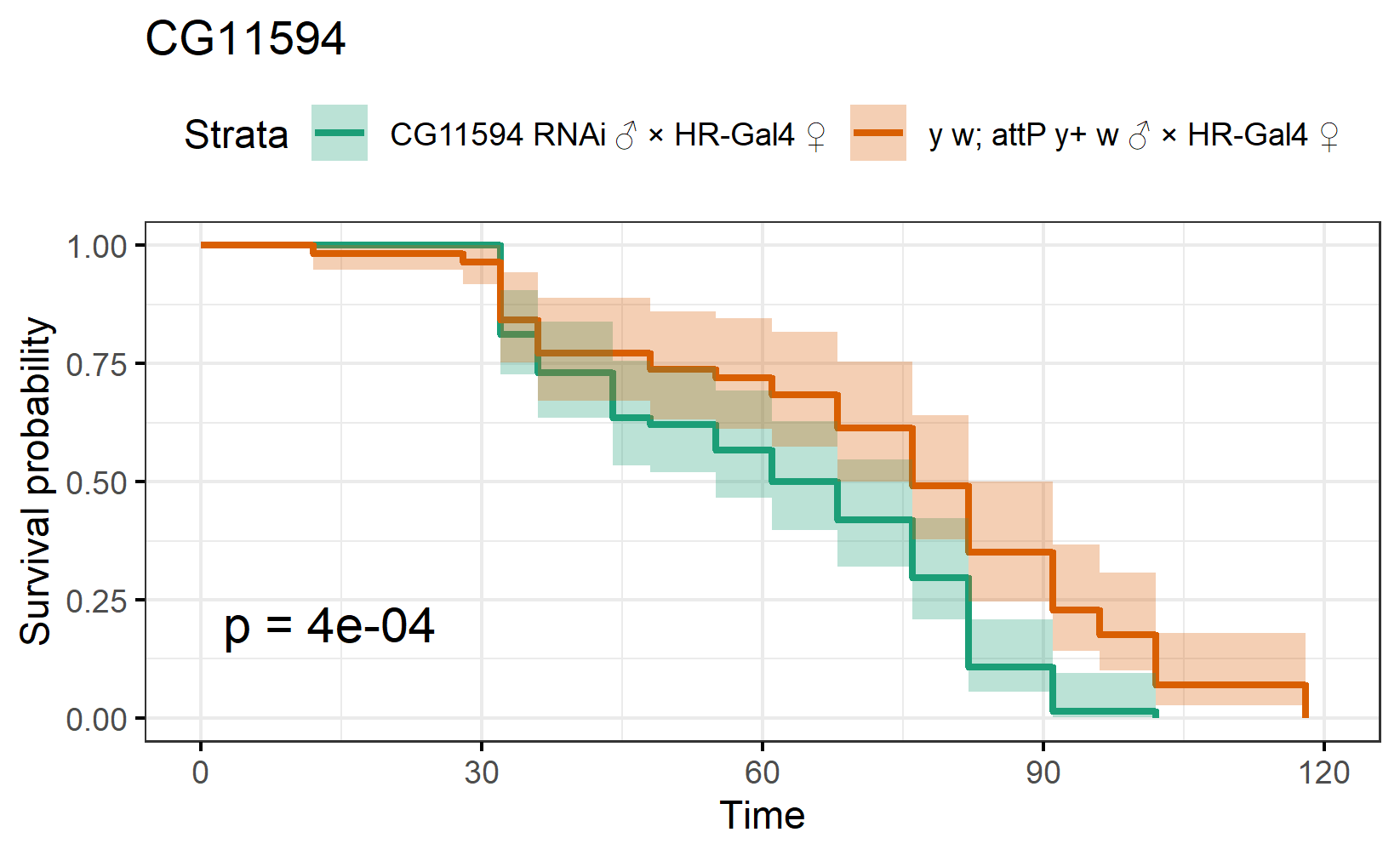

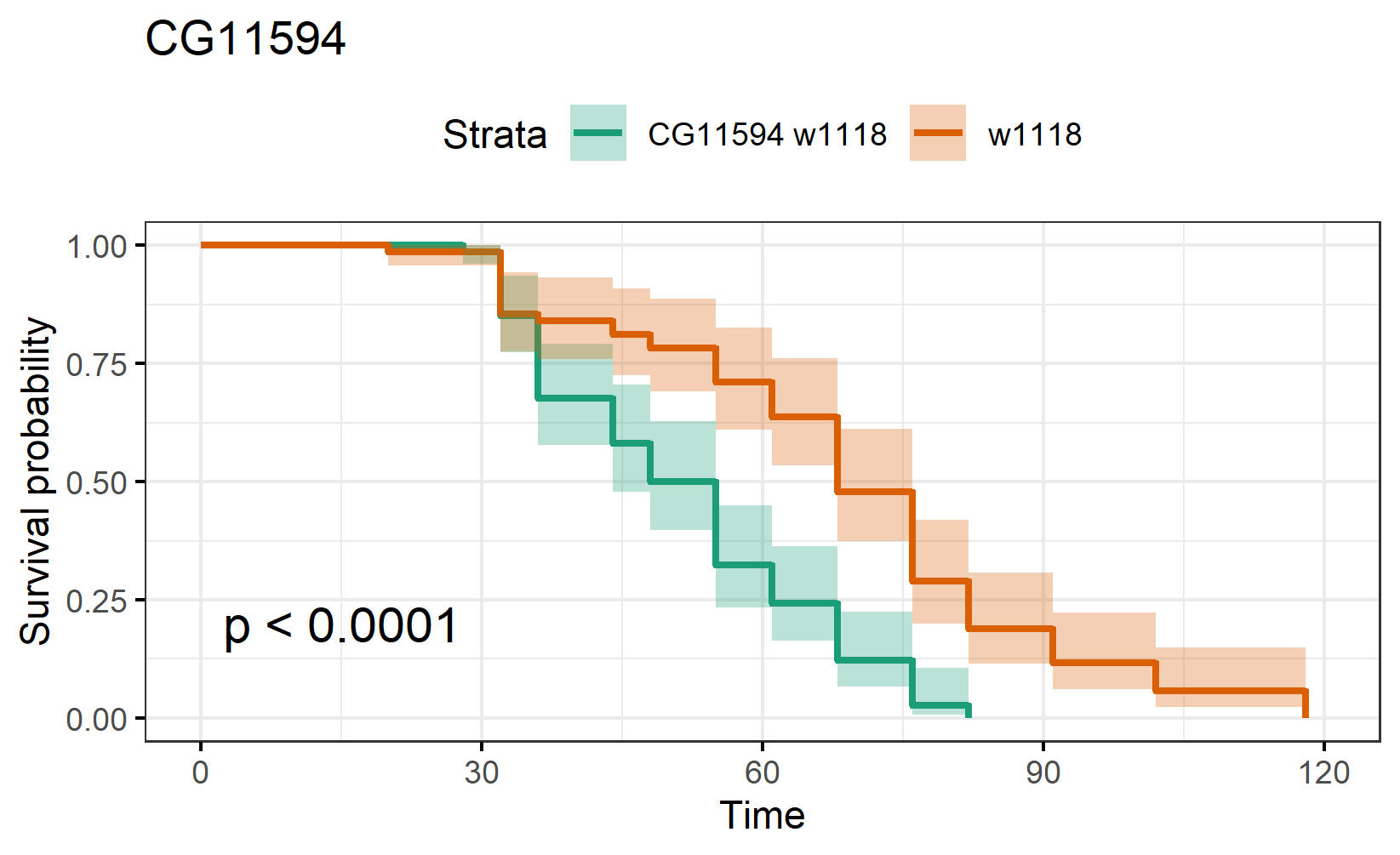

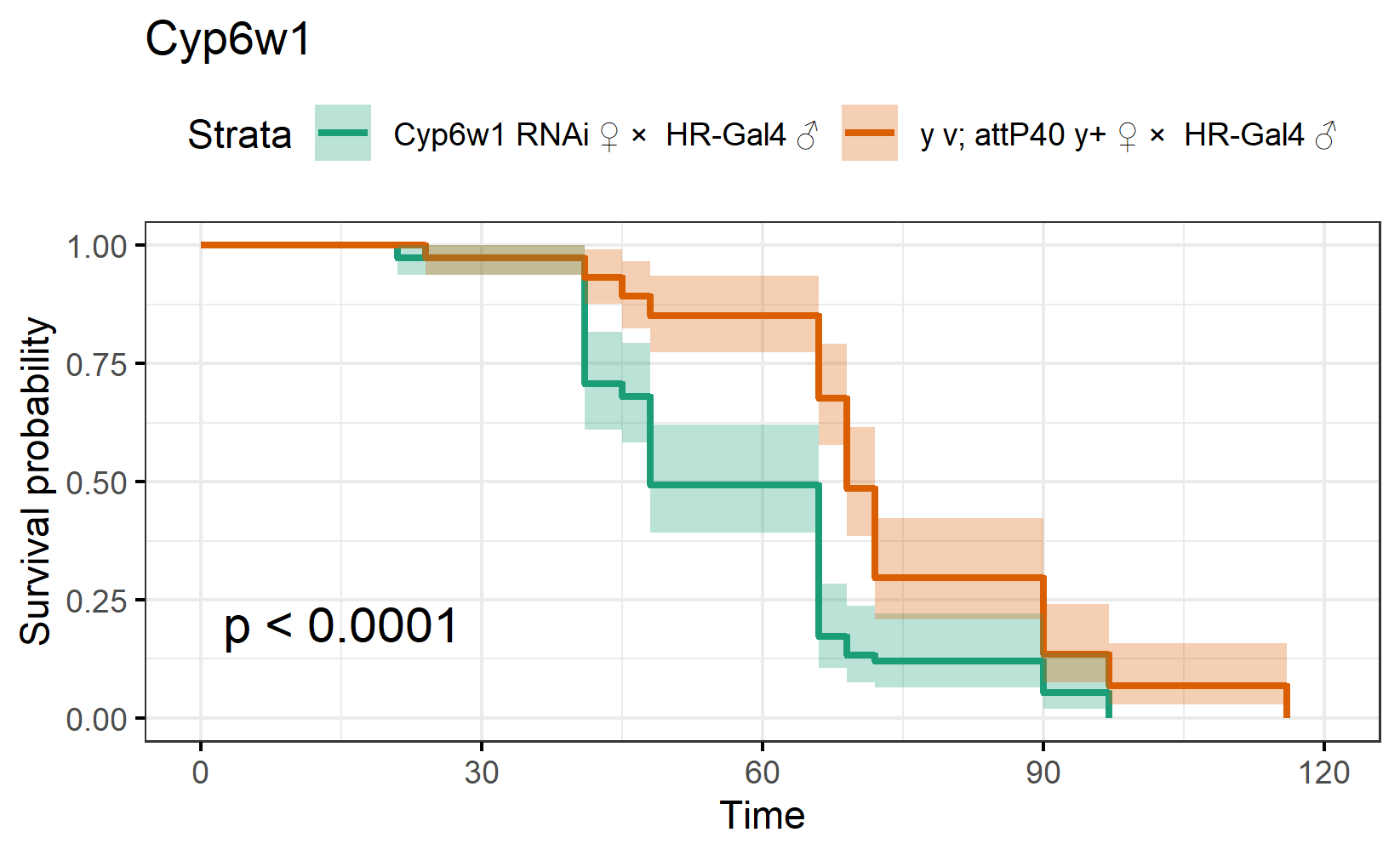

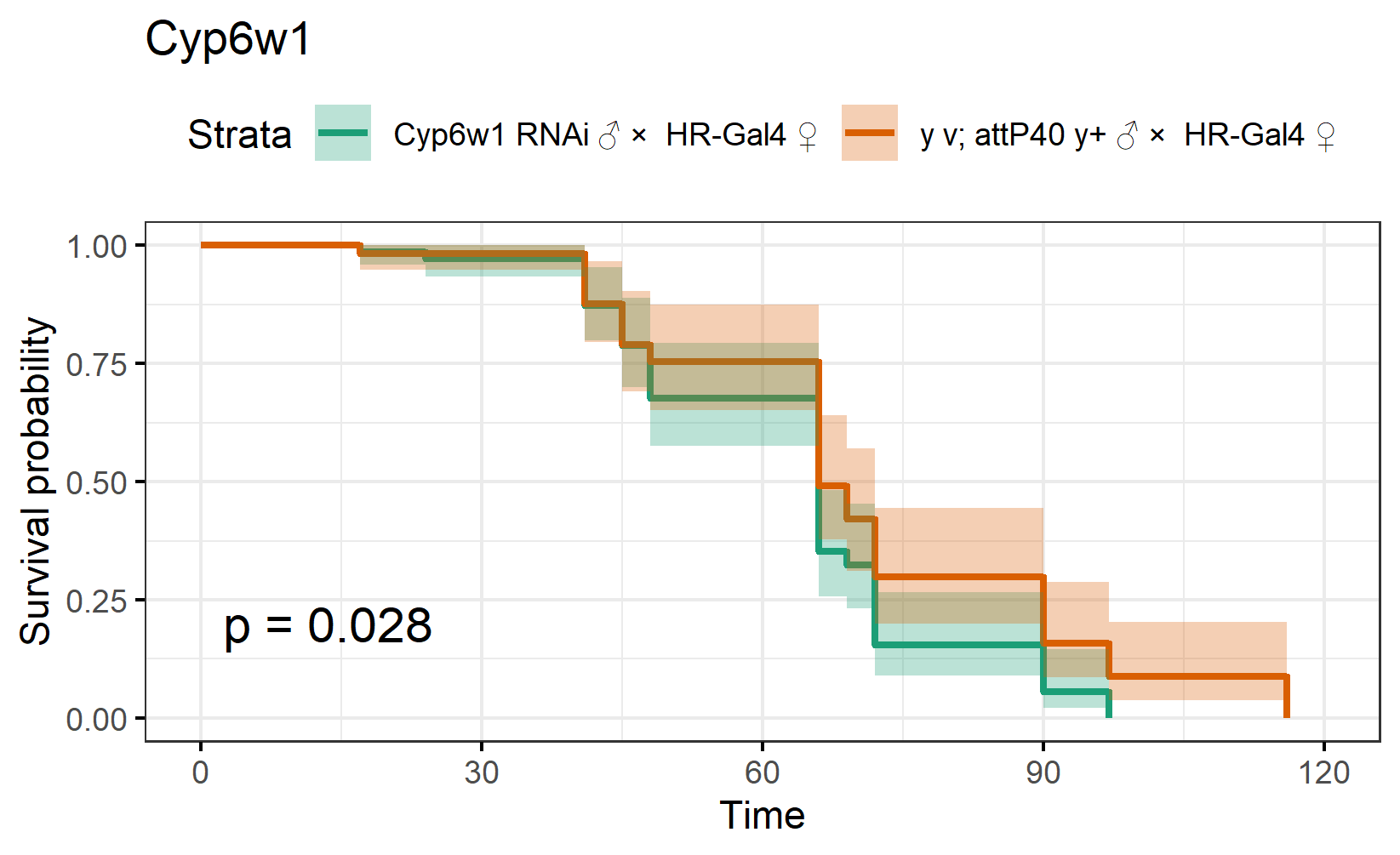

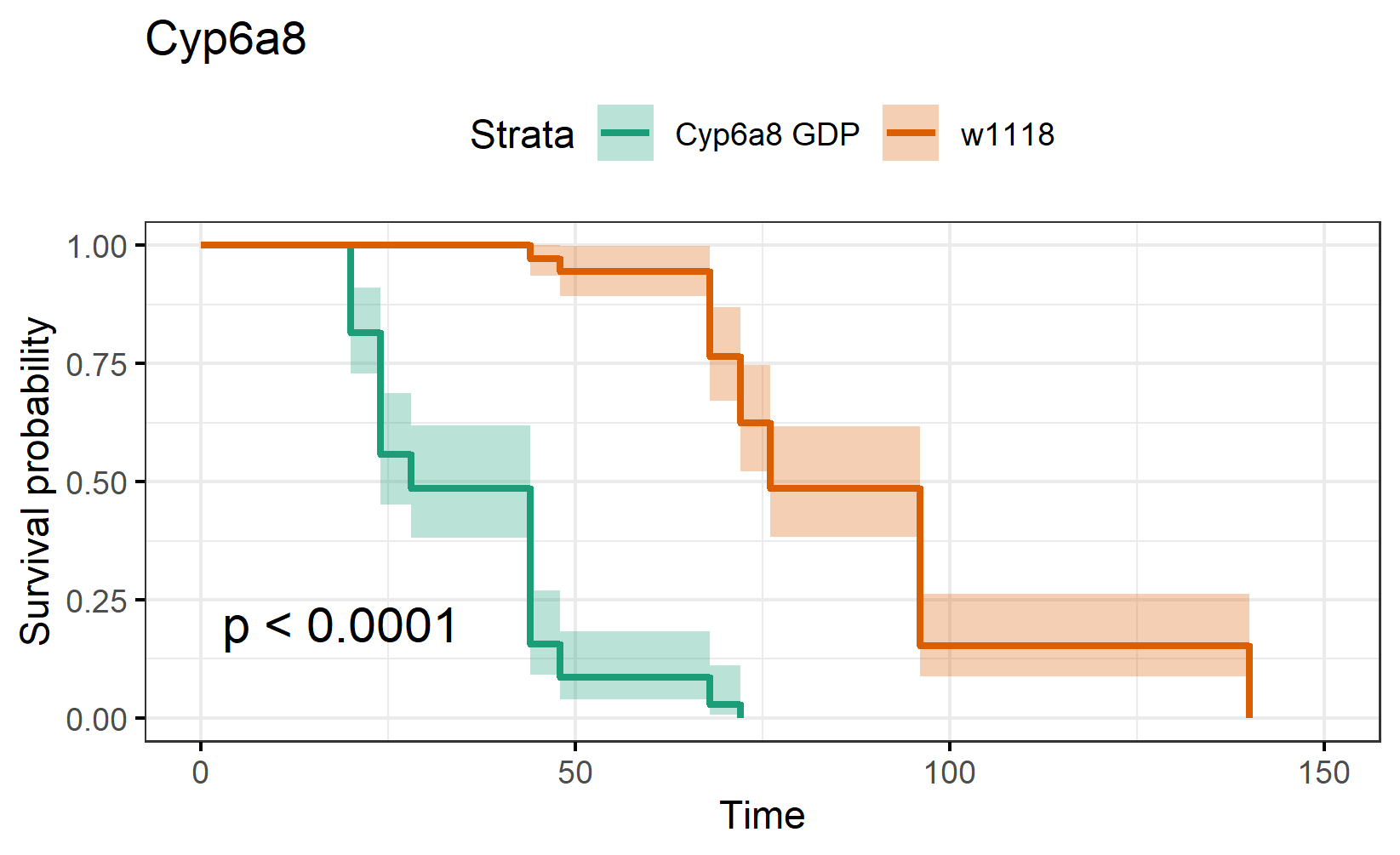

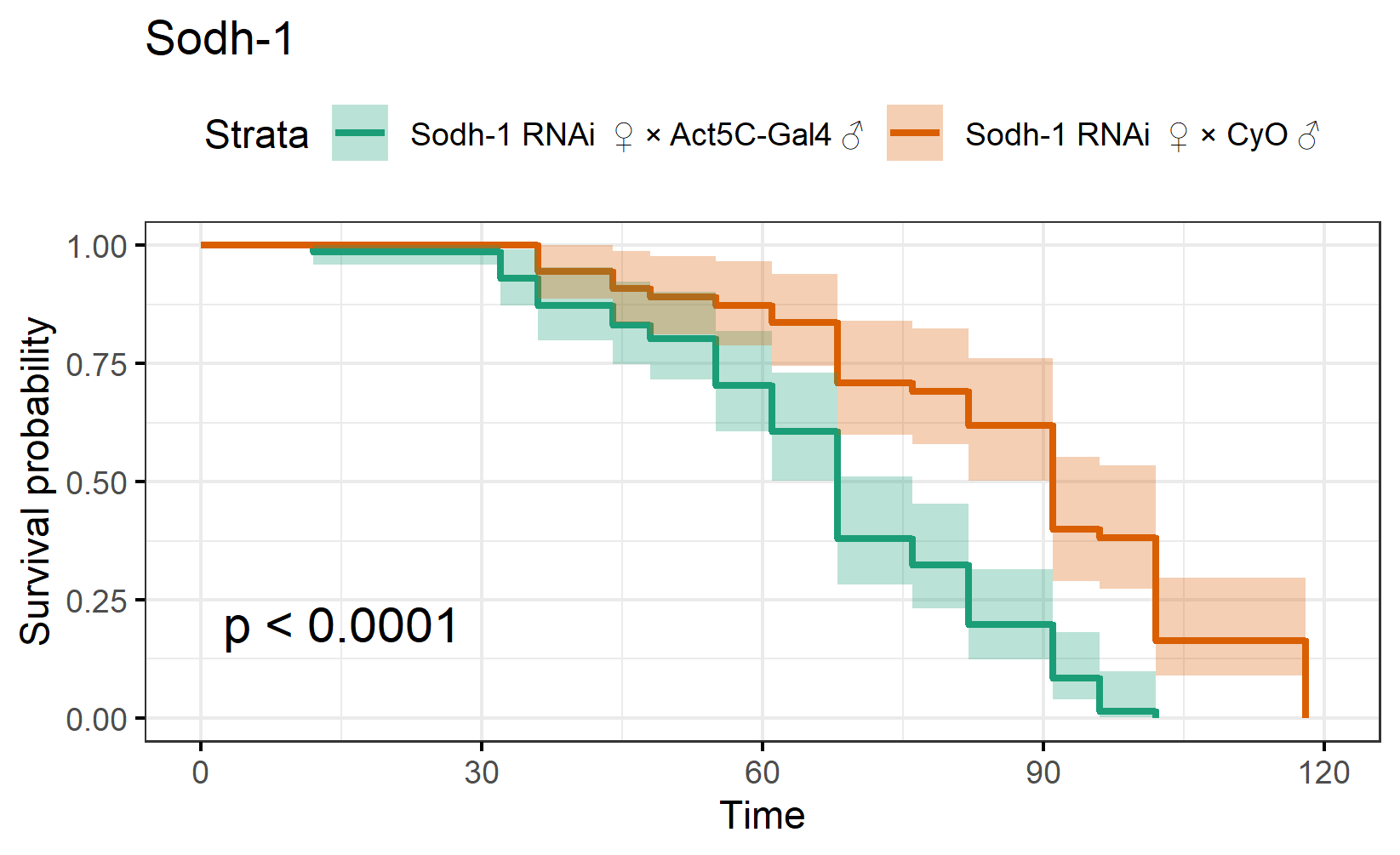

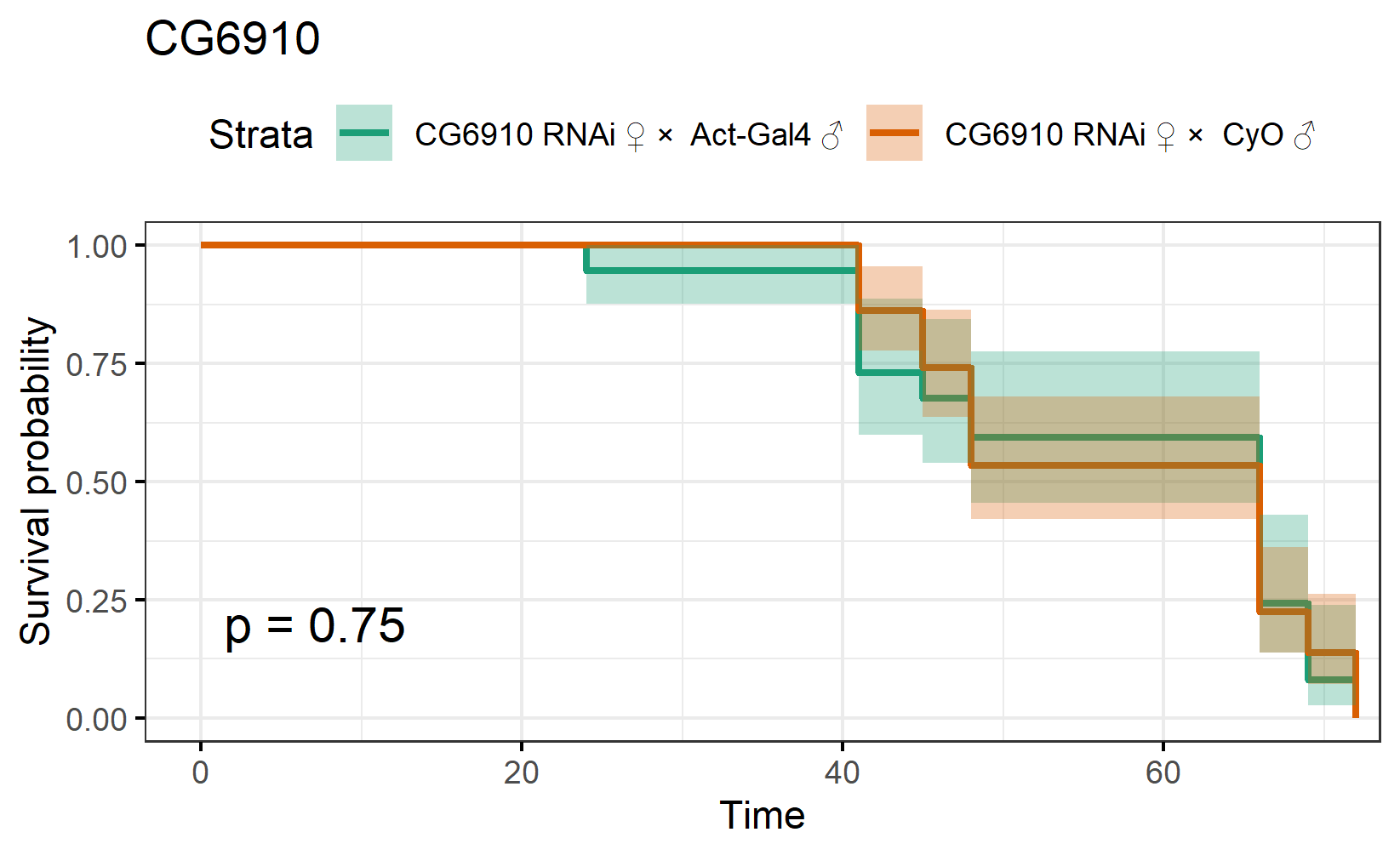

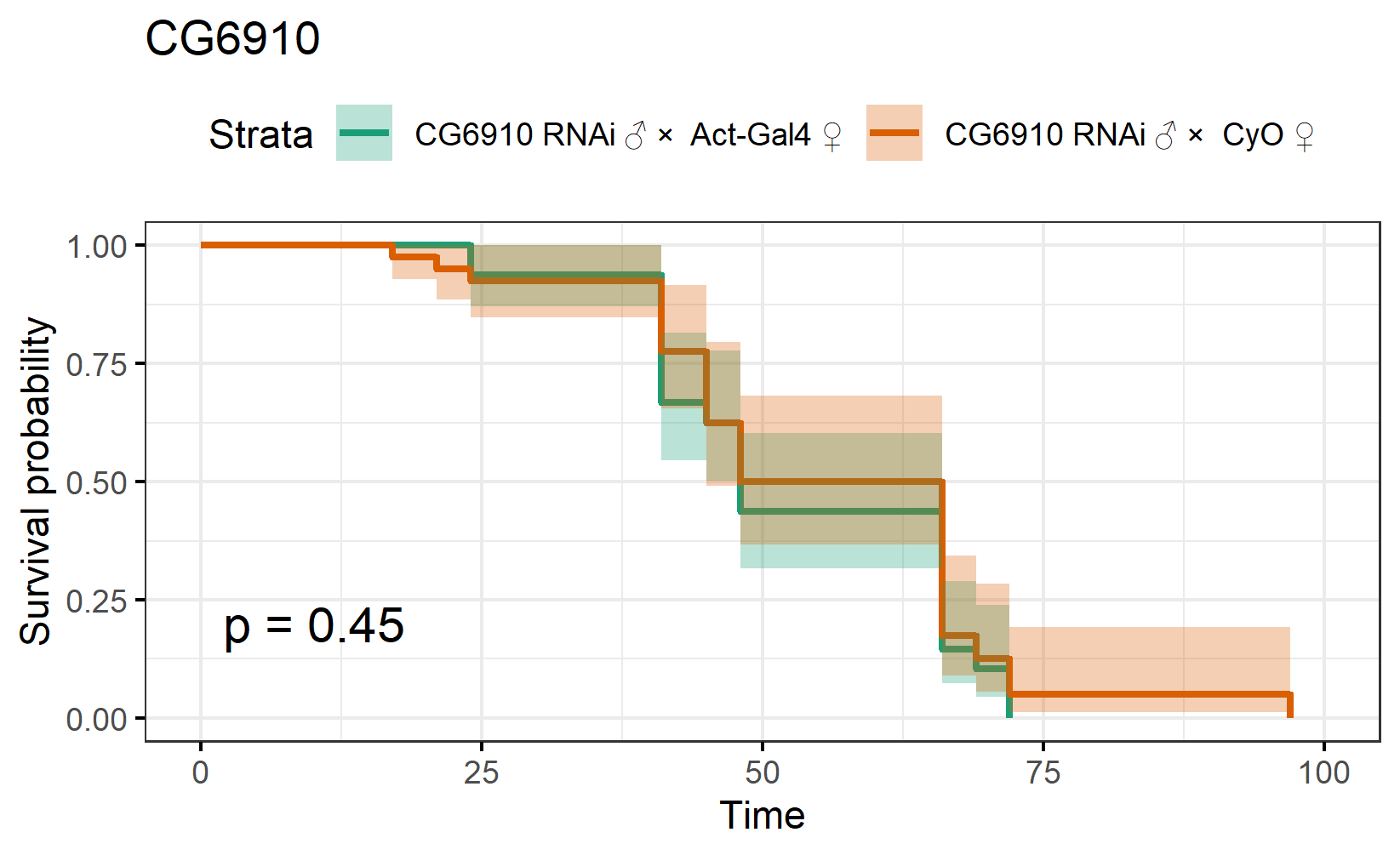

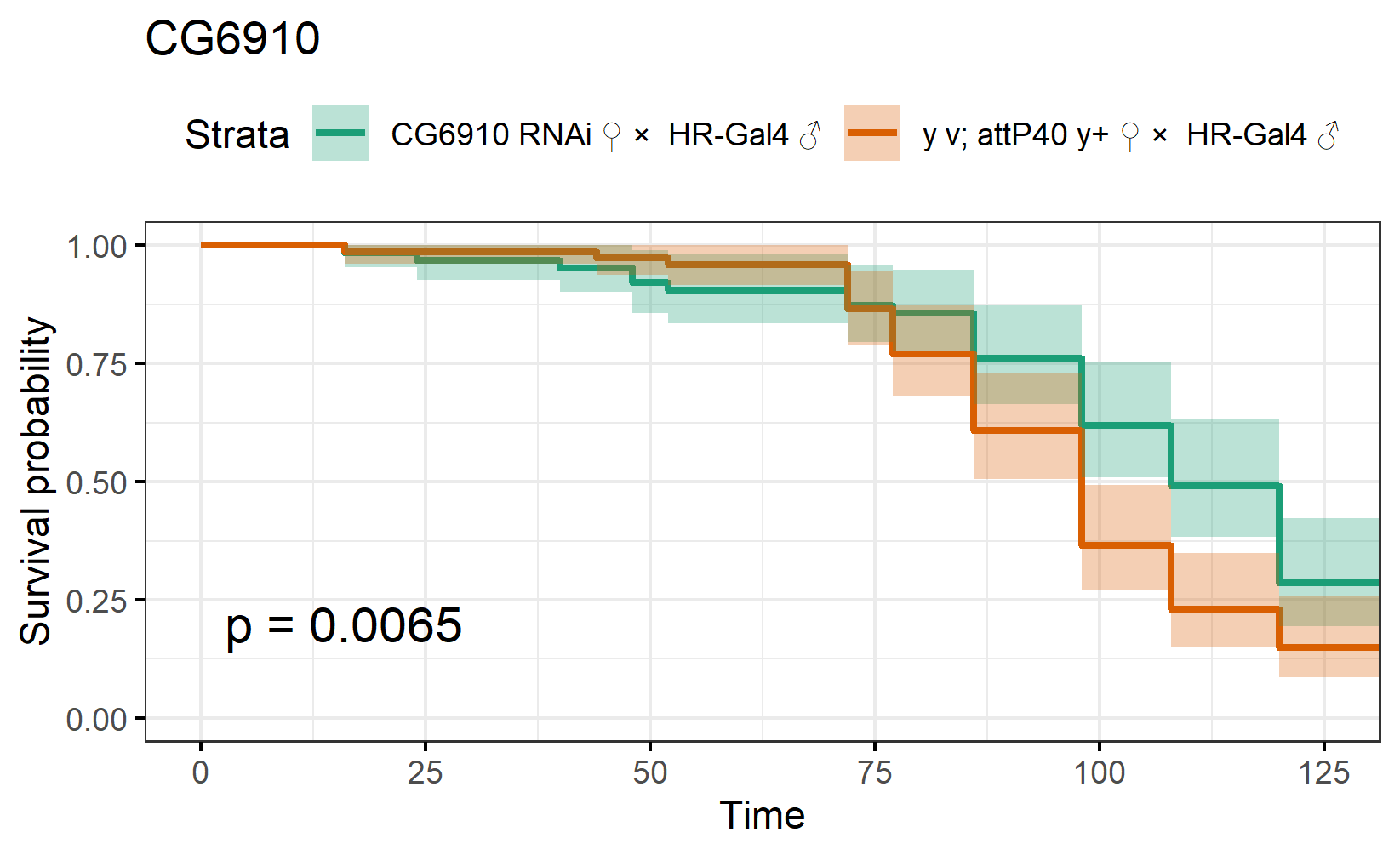

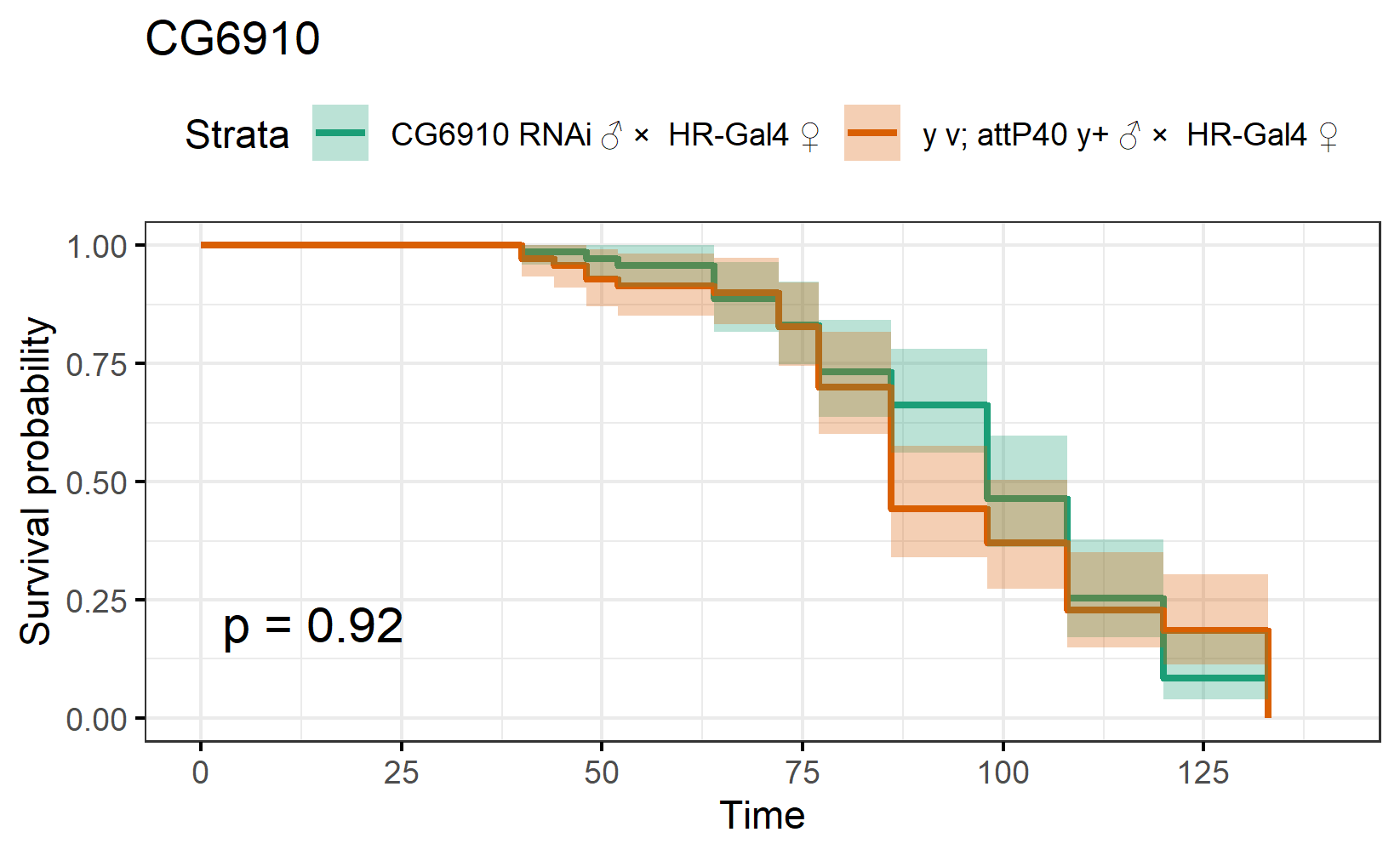

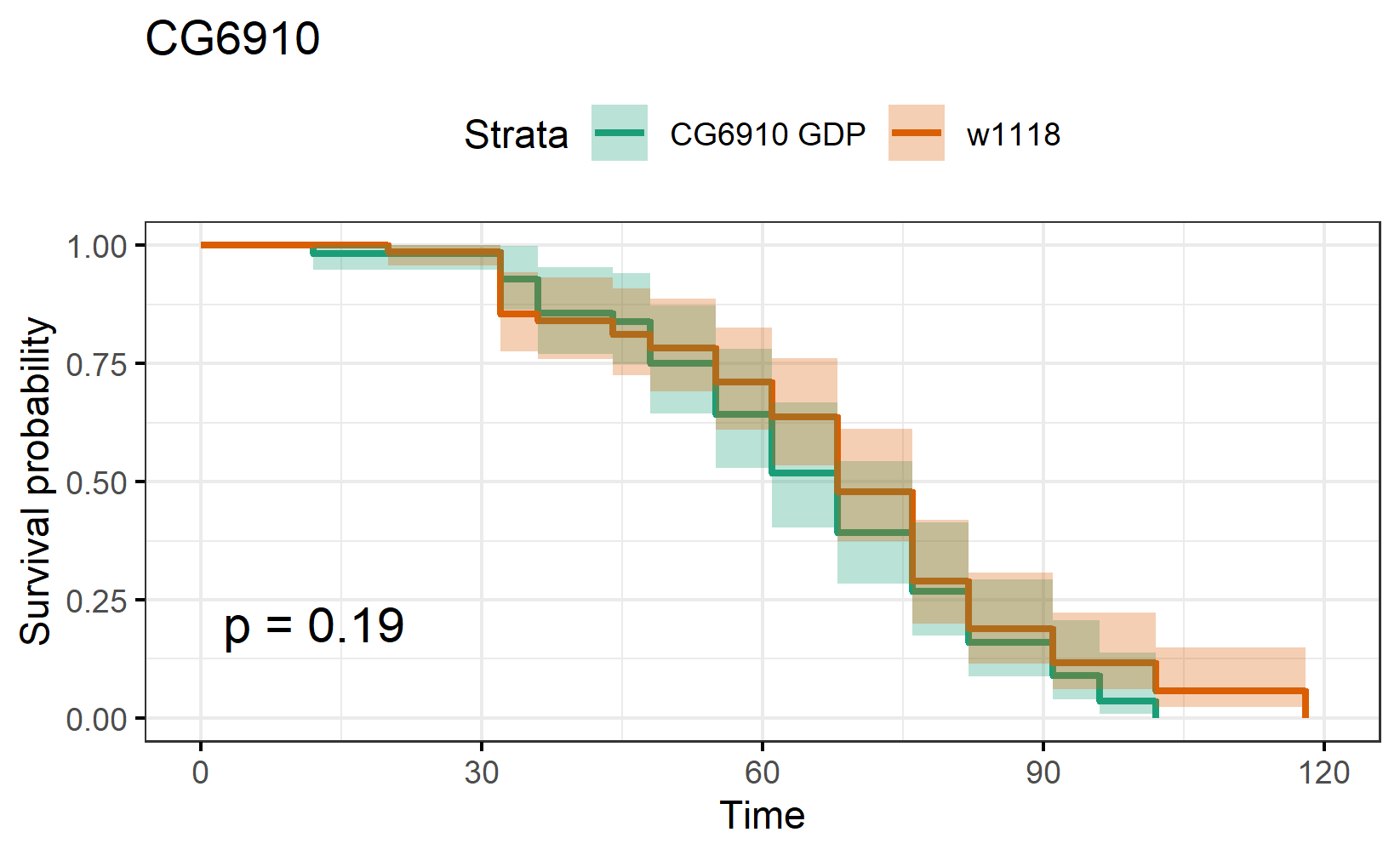

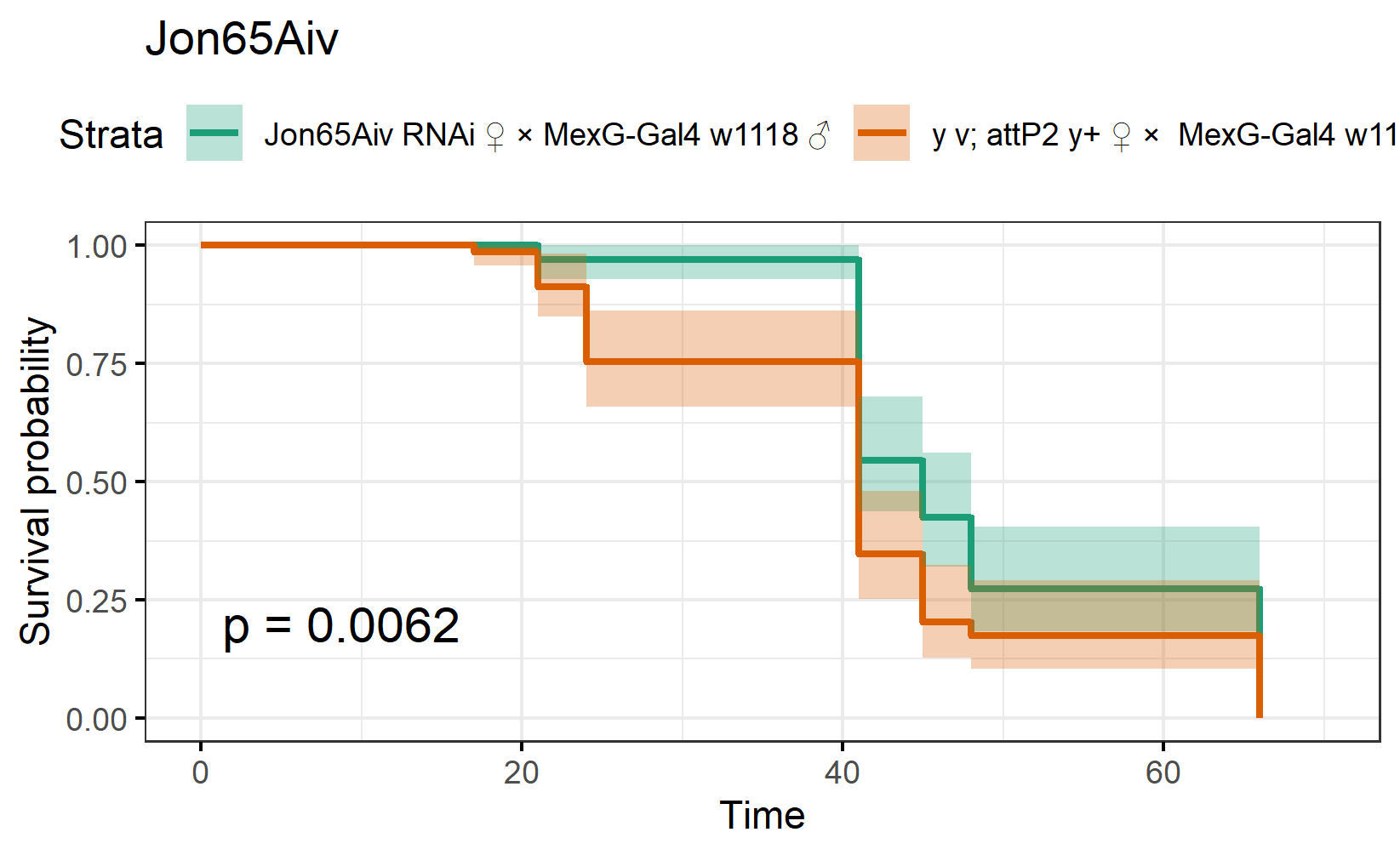

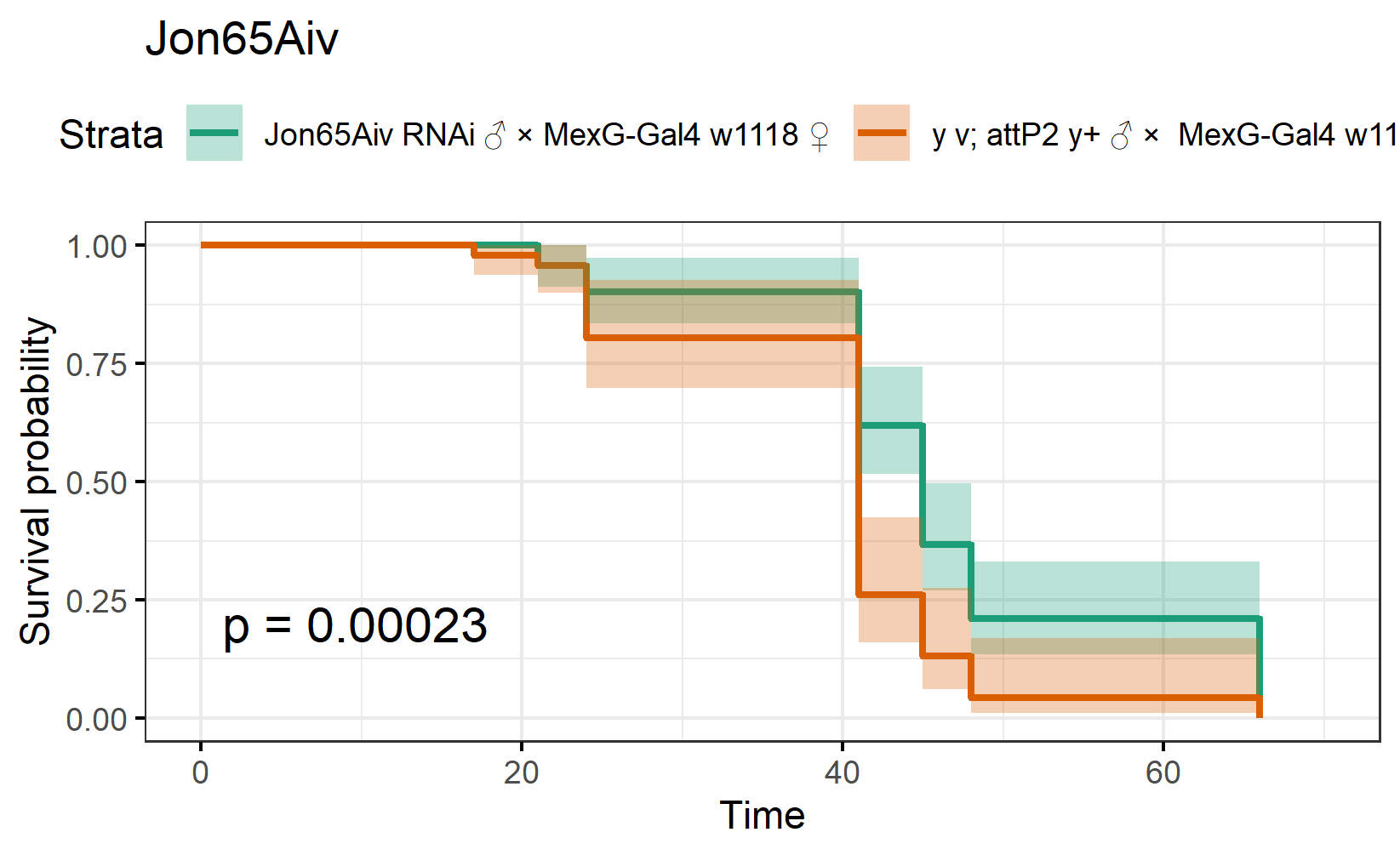

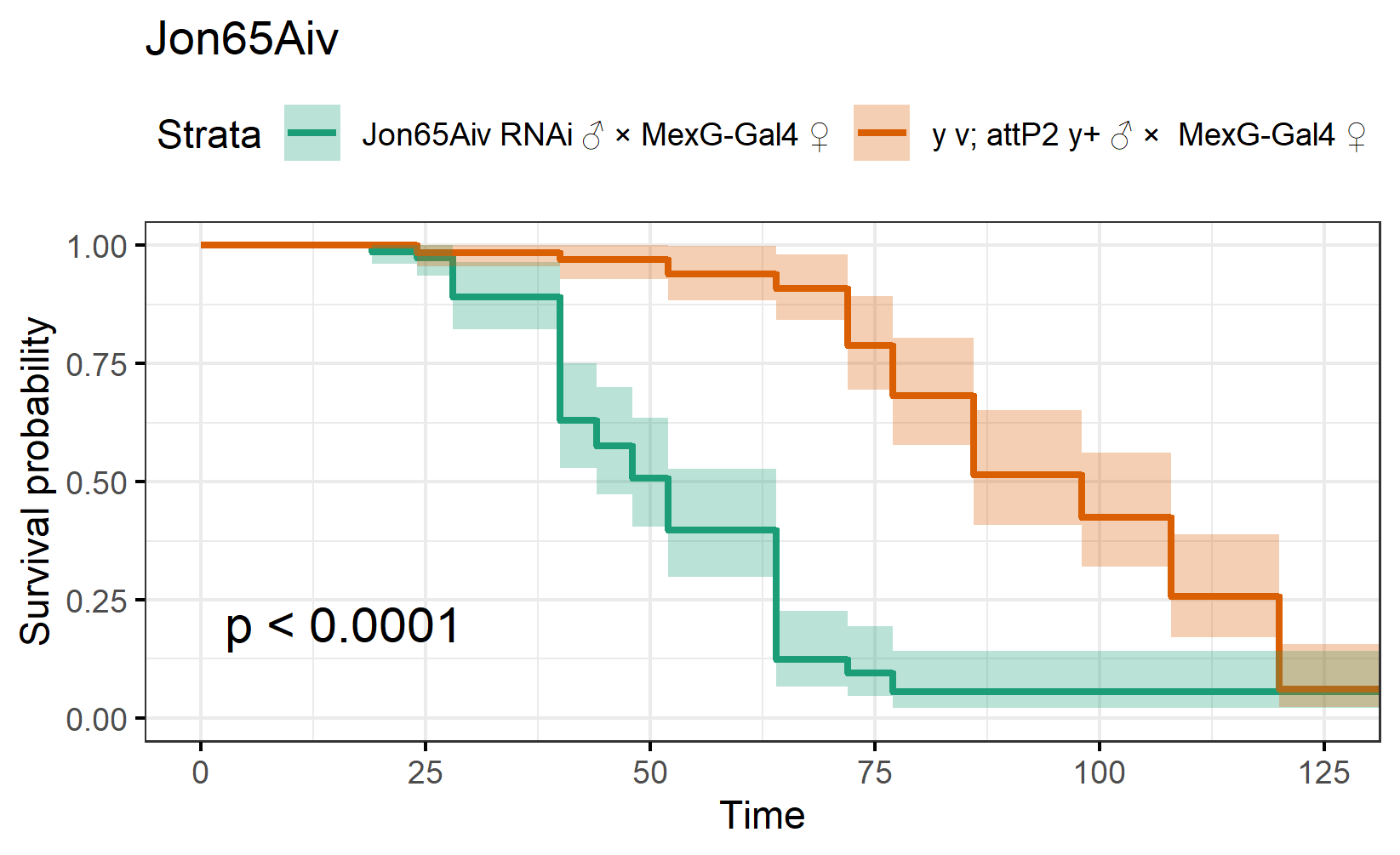

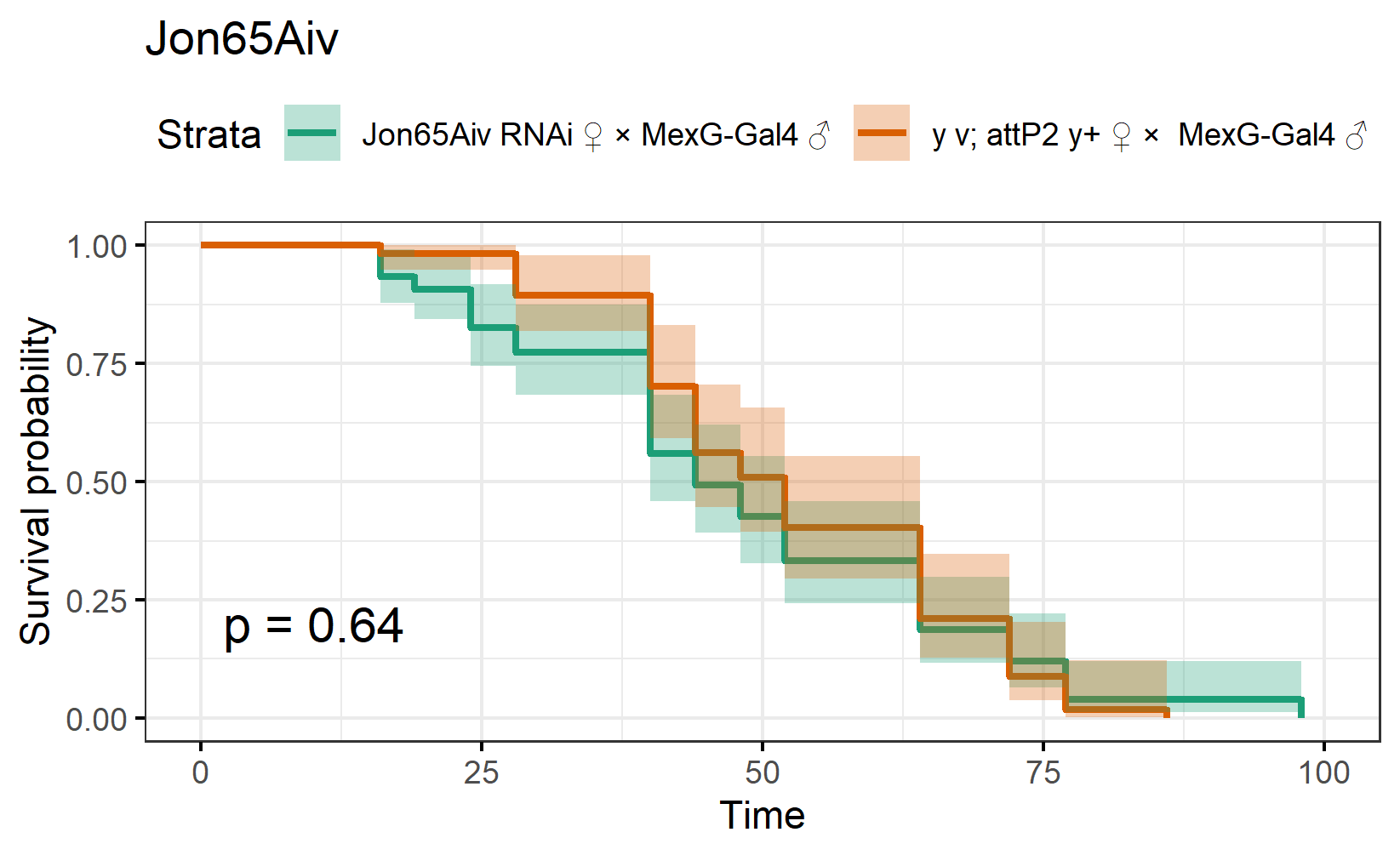

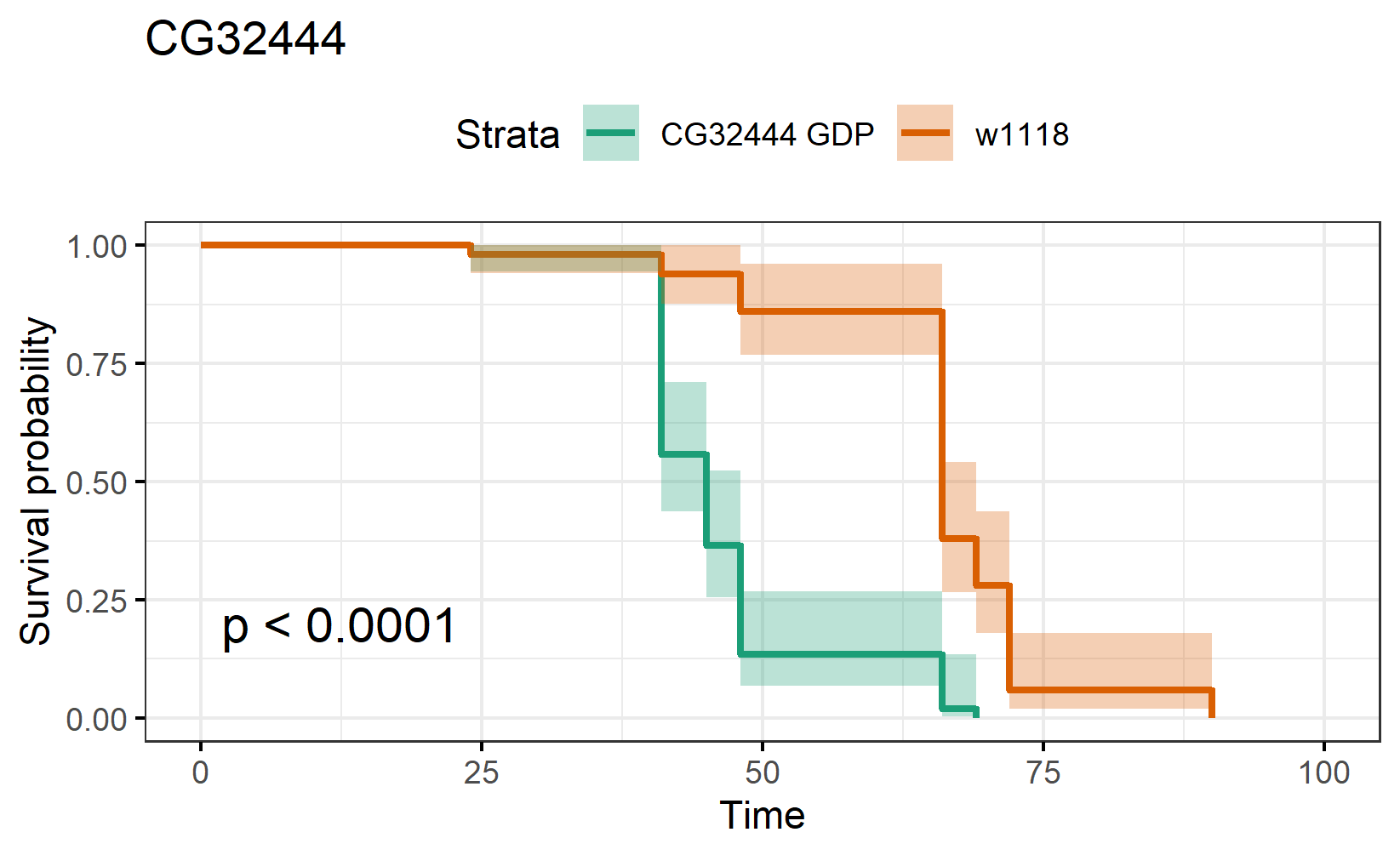

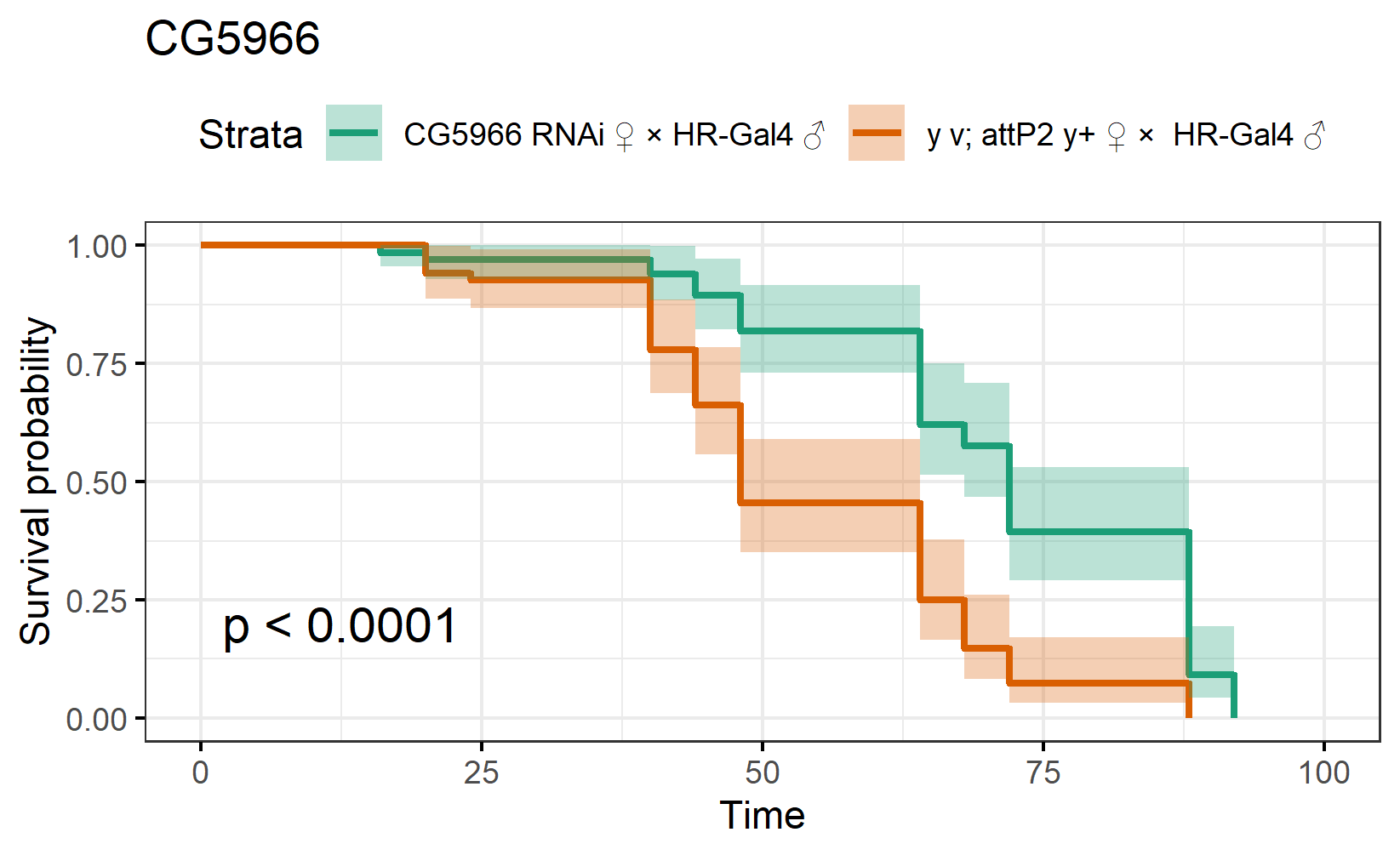

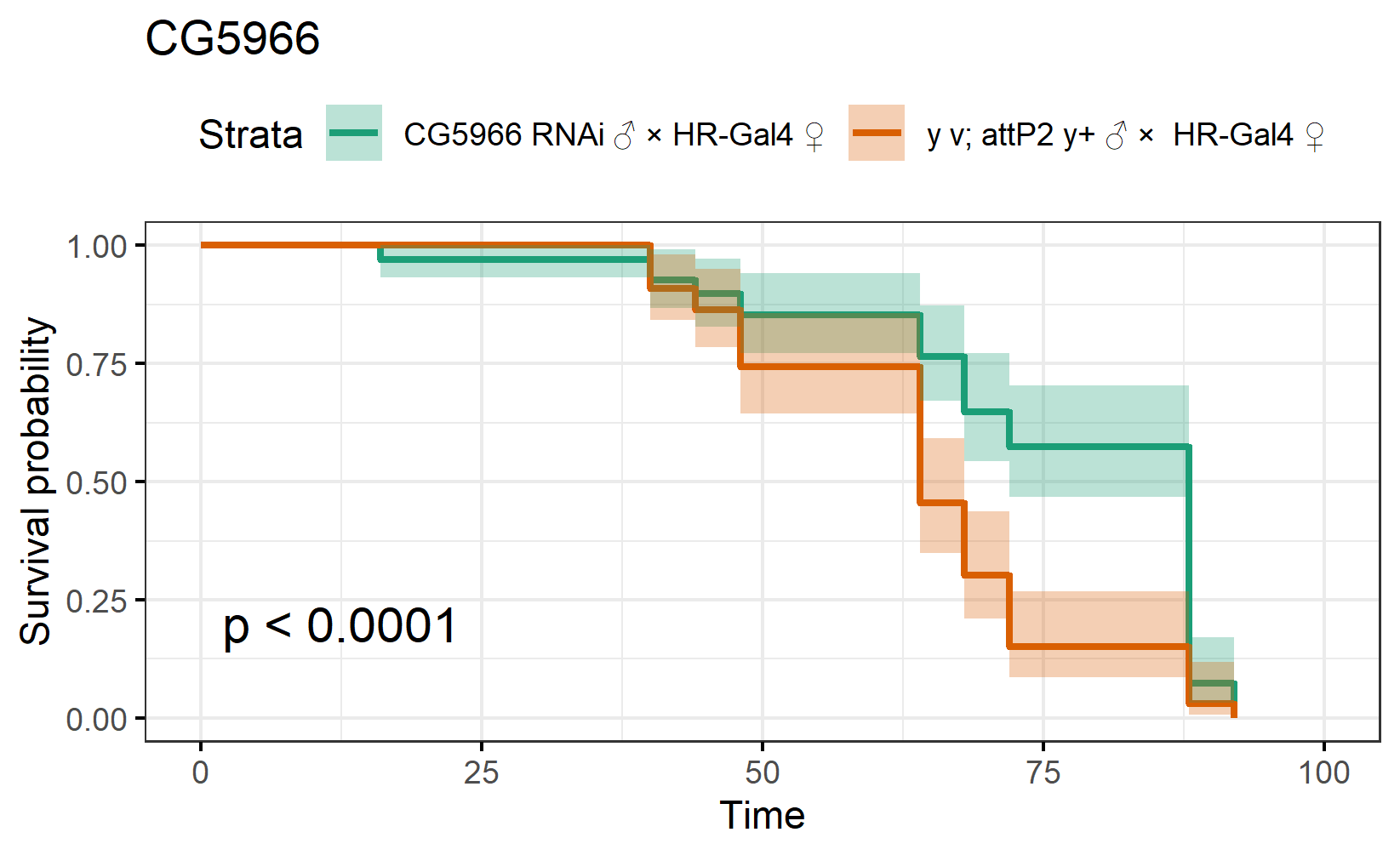

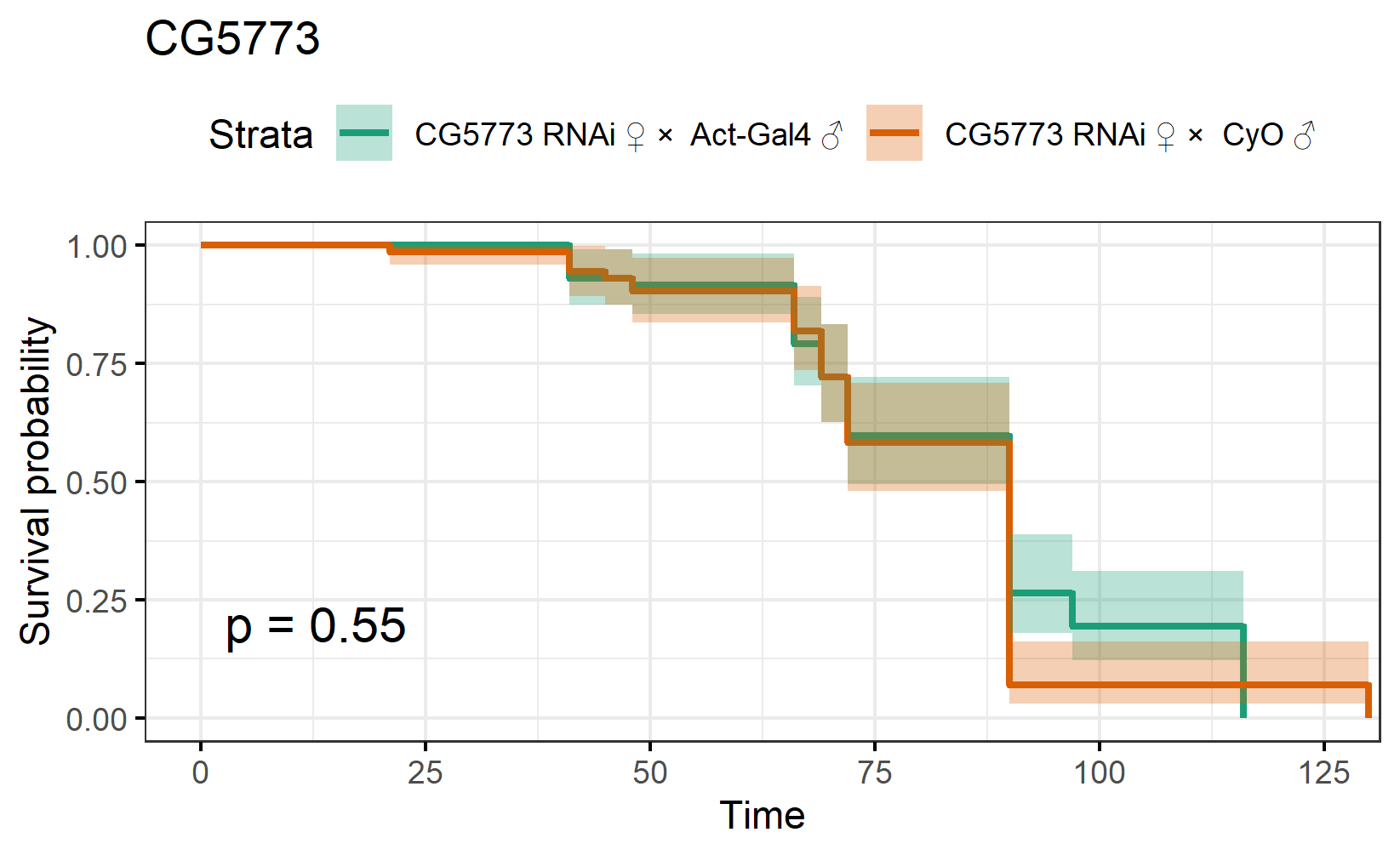

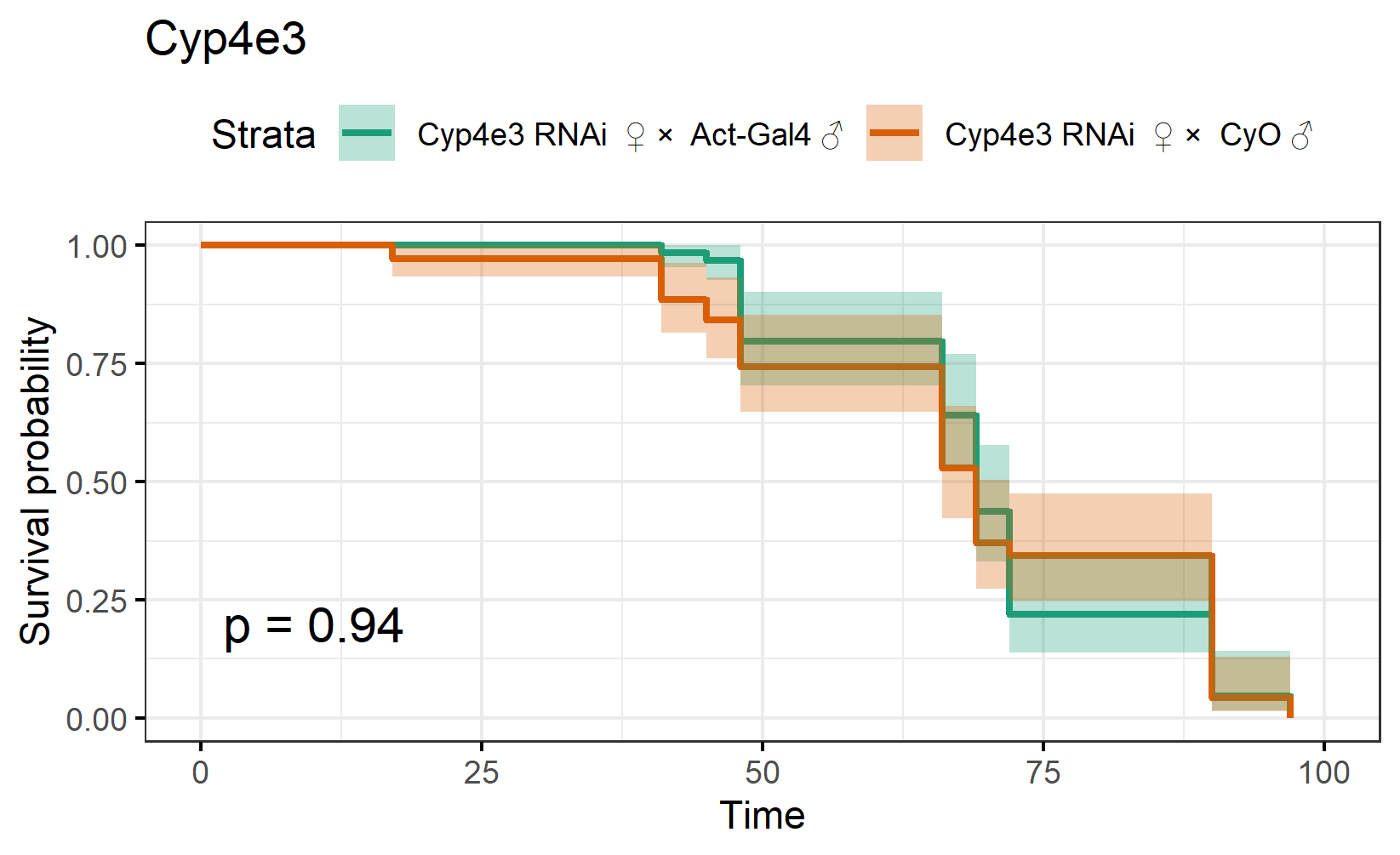

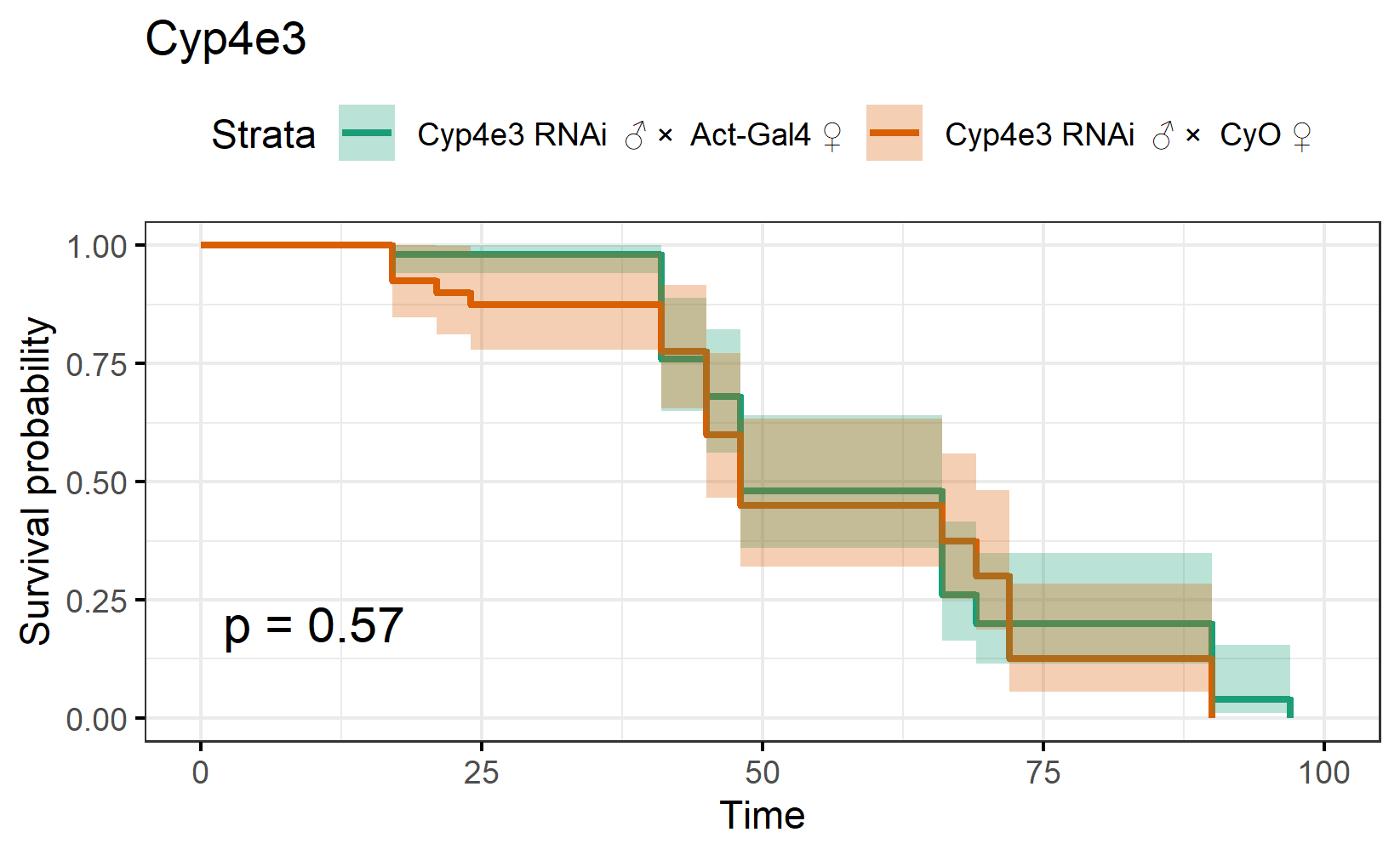

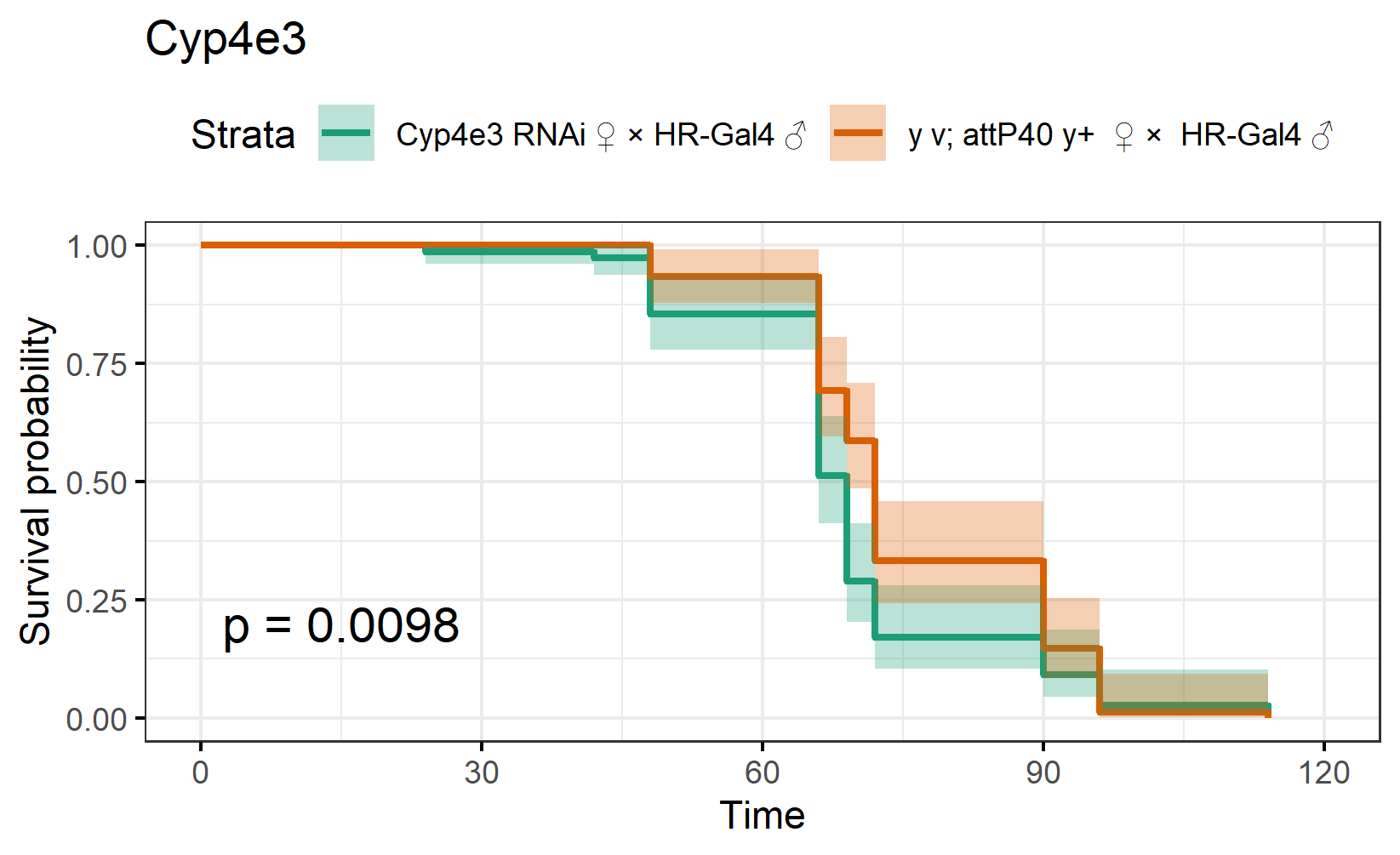

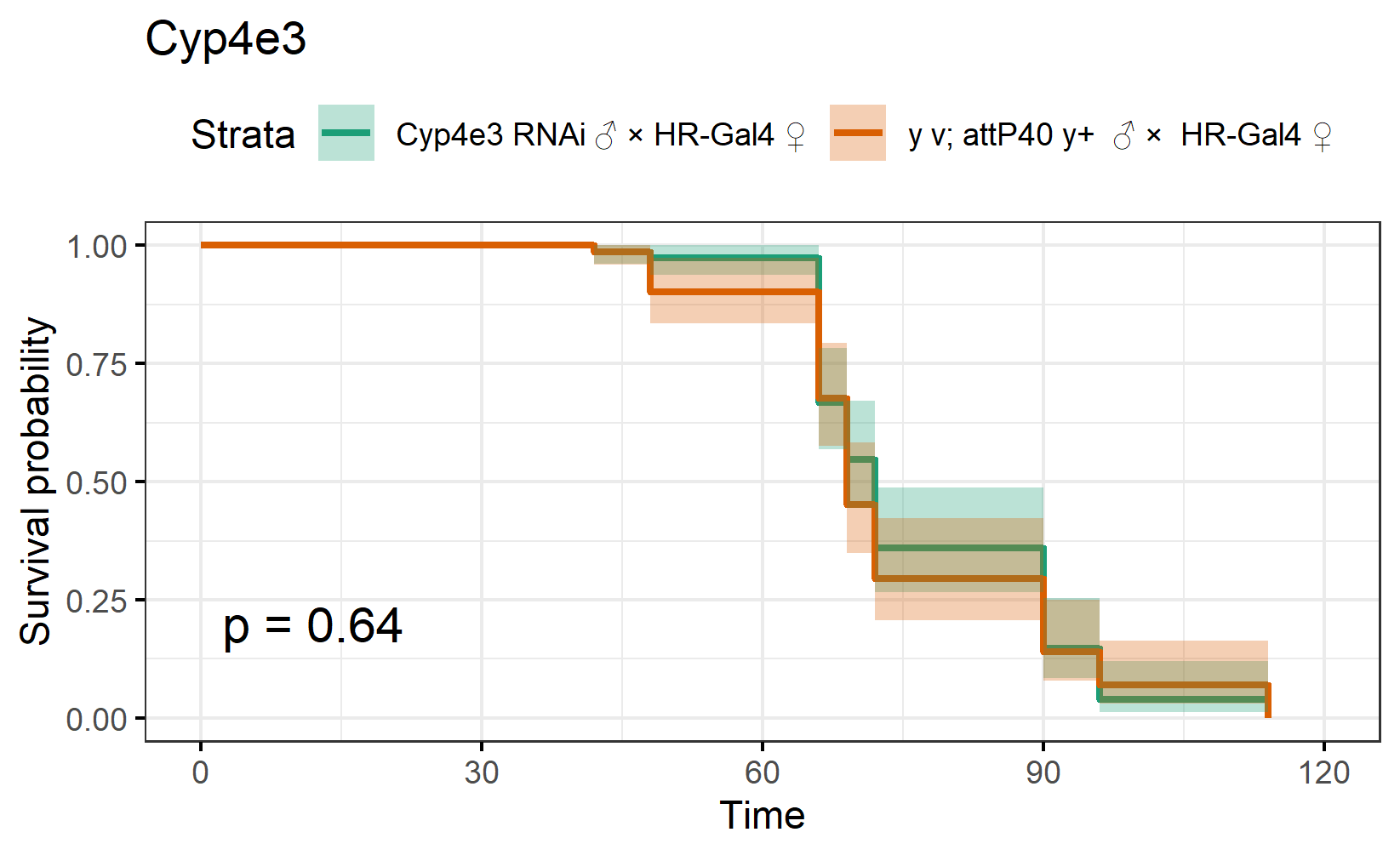

**Figure S7. Kaplan-Meier survival curves for the survival assays performed on outbred populations with and without the three candidate TE insertions.**

Shaded regions indicate the 95% confidence intervals. Statistical significance was estimated by using log-rank tests. Those plots shaded in red are the outbred populations with the TE insertion and plots in blue are the outbred populations without the TE insertion.

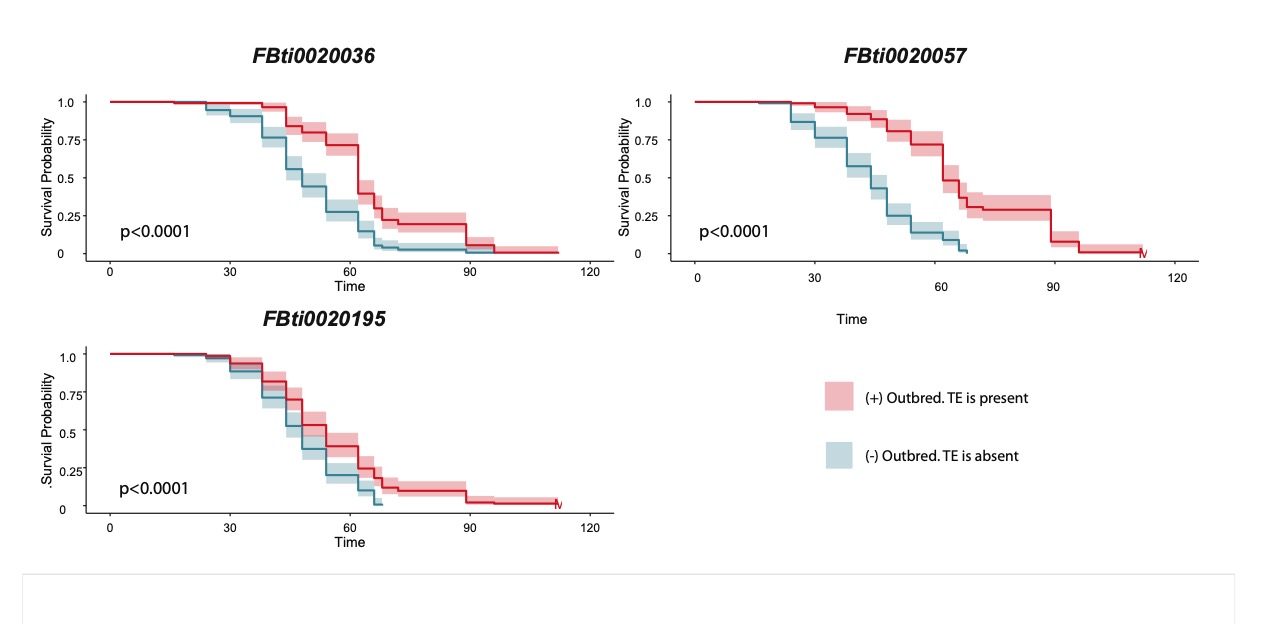
